# Reconstitution of adrenocortical functional zonation from human pluripotent stem cells

**DOI:** 10.1101/2025.05.24.655925

**Authors:** Michinori Mayama, Eoin C. Whelan, Takeshi Sato, David G. Stouffer, Adrian N. Leu, Jerome F. Strauss, Richard J. Auchus, Kotaro Sasaki

## Abstract

The adrenal cortex produces essential steroid hormones through a concentric zonal architecture, established by the centripetal trans-differentiation of subcapsular progenitors within a capsule-derived niche. To capture this complexity, we establish a human pluripotent stem cell-derived adrenal organoid system that faithfully recapitulates this process. RSPO3/WNT signaling from the capsule specifies definitive zone (DZ) progenitors from the adrenal primordium, which then differentiate into a cortisol-producing transitional zone and an androgen-producing fetal zone under the influence of RSPO3 and ACTH. Loss of *NR0B1* impairs DZ specification and triggers direct adrenal primordium-to-fetal zone conversion, mirroring the mechanism of X-linked adrenal hypoplasia congenita. When DZ cells are encapsulated with capsule cells separately derived from pluripotent stem cells, they reconstitute zonation in vivo, forming ACTH-responsive tissue that produces both cortisol and androgens. This organoid platform offers a powerful tool to dissect human adrenal development and establishes a foundation for regenerative therapies targeting adrenal diseases.

## Introduction

The adrenal cortex is the major endocrine hub for synthesis of steroid hormones essential for body homeostasis. The adrenal cortex is comprised of three functionally distinct zonal structures: the outermost mineralocorticoid-producing zona glomerulosa (zG) essential for body fluid homeostasis, the intermediate glucocorticoid-producing zona fasciculata (zF) critical for stress response, glucose homeostasis and immune regulation, and the innermost androgen-producing zona reticularis (zR) involved in sex organ development and adrenarche ^1^. The definitive zone (DZ), also known as the permanent cortex, represents the outermost prenatal cortex and appear to bear subcapsular progenitor cells responsible for the formation of the adult cortex ^2–5^. Postnatally, these cells give rise to the zG, from which the remaining adult adrenal cortex (i.e., zF and zR) is generated through continuous centripetal “trans-differentiation” ^6,7^. The functional modularity of the adrenal endocrine system is ensured by the formation of these distinct zones.

In humans, development of functional and structural cortical zonation first occurs in the prenatal period (i.e., the definitive zone [DZ], transitional zone [TZ] and fetal zone [FZ], corresponding to adult zG, zF and zR, respectively) ^2–4^. During this period, androgens produced by the FZ promote maturation of various organs and are integral for the function of the feto-placental unit, which plays a role in pregnancy maintenance ^2,8^. TZ-derived cortisol provides negative feedback to adrenocorticotropic hormone (ACTH) released from the pituitary gland and its loss in primary adrenal insufficiency results in ACTH-driven androgen excess from the FZ, thereby virilizing female fetuses ^1,9^. Postnatally, patients with primary adrenal insufficiency develop life-threatening acute adrenal failure. Thus, the developmental origin and molecular underpinning of human adrenocortical transdifferentiation in both the prenatal and postnatal periods is of importance in understanding causes of primary adrenal insufficiency and developing future regenerative therapy.

In humans, adrenal organogenesis starts with fate specification of the adrenogenic coelomic epithelium at ∼3-4 weeks post fertilization (wpf), followed by establishment of the adrenal primordium (AP) at ∼5 wpf, which subsequently culminates in formation of functionally distinct zones: the peripherally located DZ with putative progenitor potential, the intermediate cortisol-producing TZ, and the androgen-producing innermost FZ ^10–13^. In mice, the FZ and DZ originate independently from the AP, and the DZ does not appear to contribute to the FZ ^14,15^. However, our scRNA-seq data suggest that the FZ in humans can originate either directly from the AP or indirectly via the DZ, likely through trans-differentiation, akin to what is seen in adult adrenal cortex ^13^. However, such a developmental cell trajectory has not been experimentally validated, and the signaling/genetic basis of it remains largely unknown.

In mice, progenitor cells responsible for homeostatic turnover of the adrenal cortex are located in the subcapsular region and are maintained through canonical WNT/β-catenin signaling, generated in an autocrine/paracrine manner and potentiated by RSPO3 secreted from the capsule (Cap), the tissue overlying the cortex ^16–18^. Reciprocally, Cap cells are supported by SHH-Smoothened signaling provided by cortical cells ^15,19^. Furthermore, adrenocortical tissue homeostasis is maintained through antagonistic action of capsule-derived canonical WNT/β-catenin signaling and differentiation-promoting endocrine ACTH/PKA signaling ^20,21^. ACTH is also critical in human adrenal development, as evidenced by marked atrophy of the FZ in anencephalic fetuses that lack a pituitary gland, the sole organ producing ACTH ^8,22,23^. Moreover, patients bearing mutations in *WNT4*, a ligand of the canonical WNT pathway, develop severe primary adrenal insufficiency ^24^. Although these observations suggest that both WNT- and ACTH-signaling are operative in the prenatal human adrenal cortex, how these signaling pathways cooperate to establish zonal structures in prenatal adrenal cortex remains unknown.

While rodent models provide valuable insights into adrenal development and steroidogenesis, species-specific differences limit our understanding of this process in humans ^13,25^. As mechanistic evaluation in human embryos is untenable, we recently developed a human induced pluripotent stem cell (iPSC)-derived prenatal adrenal organoid system that partially recapitulates functional adrenocortical development and steroidogenesis ^26^. While these organoids contain androgen-producing FZ-like cells (FZLCs) and a small number of DZ-like cells (DZLCs), they do not contain cortisol-producing TZ-like cells (TZLCs), limiting their therapeutic use to replace cortisol-loss in patients with primary adrenal insufficiency. However, by exploiting knowledge obtained in our previously established adrenal organoid system as well as that obtained in our transcriptional analysis of prenatal human adrenal cortex in vivo, we have now successfully advanced our adrenal organoid platform. We reveal that DZLCs robustly induced by defined factors from iPSCs can reconstitute functional zonation through trans-differentiation, resulting in formation of ACTH-responsive adrenal cortex capable of producing cortisol and androgens both in vitro and in vivo. This will provide a critical foundation for deciphering the cellular and genetic basis of human adrenocortical development and steroidogenesis as well as for future cell-based therapy for primary adrenal insufficiency.

## Results

### Canonical WNT signaling provided by CapLCs is essential for derivation of DZLCs in vitro

Within human iPSC-derived adrenal organoids that consist primarily of FZLCs, we have previously identified rare NR5A1-2A-tdTomato (NT)^+^ cortical cells bearing DZ markers (e.g., MME [a.k.a. CD10]) ^26^. To improve FZLC induction efficiency, we recently modified our protocol to one in which organoids are embedded within Matrigel instead of being cultured at the air-liquid interface (fig. S1A, B) ^26^. In Matrigel organoid culture, albeit without addition of cytokines, a similar NT^+^CD10^+^ population could be identified (Fig. 1A). To identify the signaling pathways active in these cells, we FACS-sorted CD10^+^ and CD10^−^ cells among NT^+^ cells induced from iPSCs bearing NT fluorescence reporter alleles (NT 1390G3-2125, herein designated as 2125 iPSCs) and performed bulk RNA-seq analyses (Fig. 1A). This analysis revealed that CD10^+^ NT^+^ cells upregulated genes associated with canonical WNT signaling (e.g., *LEF1*, *AXIN2*) along with previously identified DZ markers (e.g., *HOPX*, *TSPAN13*, *MME*). Thus, we designated this population as DZLCs. Consistently, organoids cultured in the presence of an inhibitor of canonical WNT signaling (IWR1) or neutralizing antibody for RSPO3 (a potent activator of canonical WNT signaling) showed a significant reduction in the number/proportion of DZLCs in organoids (Fig. 1B, C). Furthermore, DZLCs were completely missing in organoids induced from bialleleic *RSPO3* mutant iPSCs, suggesting a critical role for endogenous RSPO3-mediated WNT activation in DZLC induction (Fig. 1D, fig. S1C, D).

**Fig. 1.**
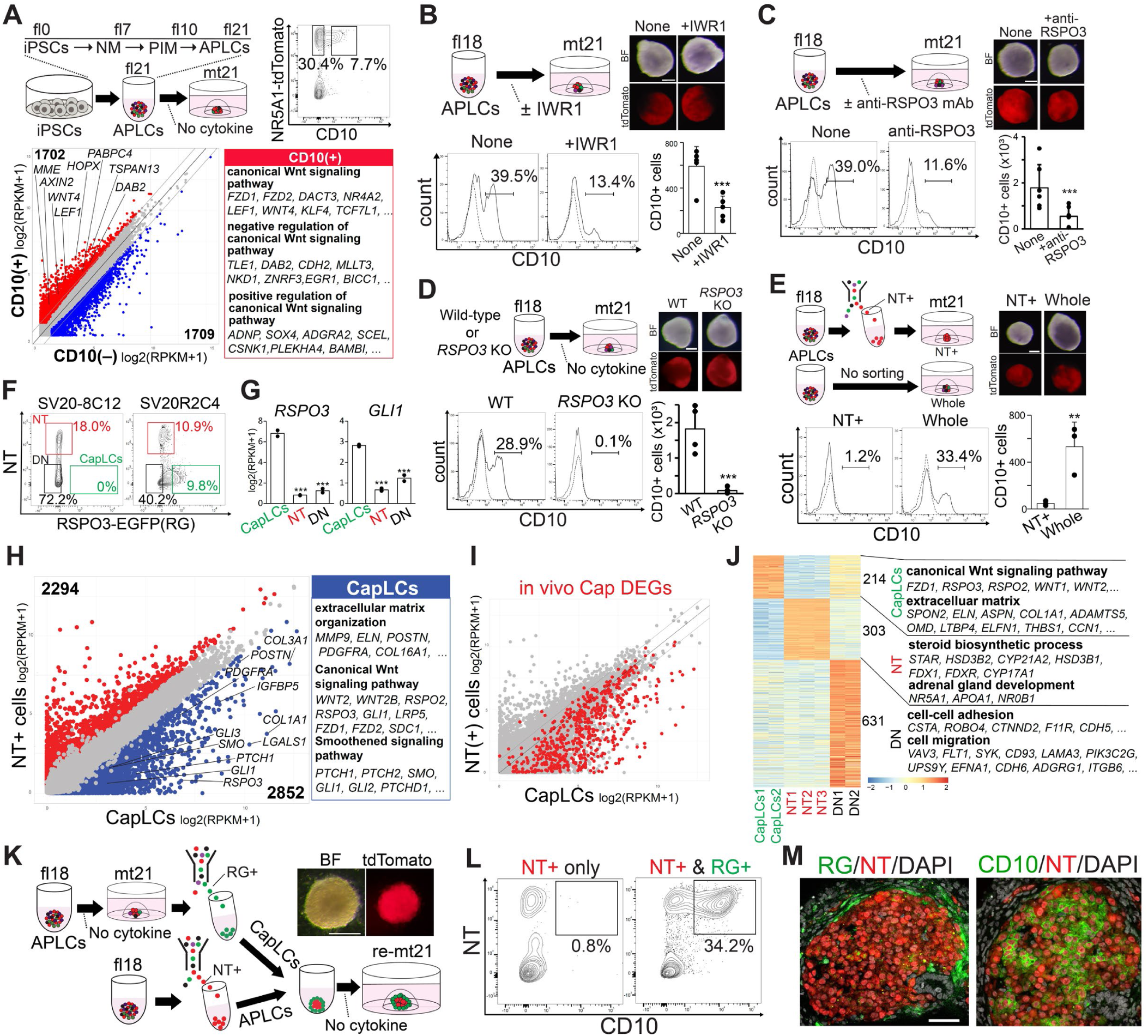
Canonical WNT signaling provided by CapLCs is essential for derivation of DZLCs in vitro. **(A)** Scheme of adrenal organoid induction. Human iPSC-derived adrenal organoids were embedded in Matrigel and cultured in base medium for 21 days (mt21) following an initial 21-day floating culture (fl21) (top left). Flow cytometry (FACS) gating strategy used to isolate NR5A1-tdTomato (NT)^+^CD10^+^ and NT^+^CD10^−^ cells at mt21 for bulk transcriptome analysis (top right). Scatter plot comparing average gene expression between NT^+^CD10^+^ and NT^+^CD10^−^ cells (bottom). Differentially expressed genes (DEGs) (log-fold change >1, FDR <0.05, log2[RPKM+1] >1) are highlighted. Representative DEGs and their associated Gene Ontology (GO) enrichments are shown on the right. NM, nascent mesoderm; PIM, posterior intermediate mesoderm; APLCs, adrenal primordium-like cells. **(B)** Organoids derived from 2125 iPSCs cultured with or without IWR1(1uM) (top left). Bright field (BF) and fluorescence images of organoids (top right). FACS histograms showing the percentage of CD10^+^ cells within the NT^+^ fraction (bottom left) and the total number of NT^+^CD10^+^ cells per organoid (bottom right). Bar graph represents mean + standard deviation (n = 5). Scale bar, 200 μm; *p < 0.05, **p < 0.01, ***p < 0.001. **(C)** Organoids derived from 2125 iPSCs cultured with or without neutralizing antibody for RSPO3 (2uM). Bar graph represents mean + standard deviation (n = 6). Scale bar, 200 μm; *p<0.05, **p<0.01, ***p<0.001. **(D)** Organoids derived from 232B6 (*RSPO3* KO) or 2125 iPSCs (parental line of 232B6), cultured in base medium only as in (A). Bar graph represents mean + standard deviation (n = 4). Scale bar, 200 μm; *p<0.05, **p<0.01, ***p<0.001. **(E)** Organoids cultured until mt21 with or without FACS-sorting of NT^+^ adrenal primordium-like cells (APLCs) at fl18. Organoids were derived from 2125 iPSCs. Bar graph represents mean + standard deviation (n = 3). Scale bar, 200 μm; *p < 0.05, **p < 0.01, ***p < 0.001. **(F)** FACS gating strategy for organoids at mt21 from SV20R2C4 iPSCs bearing *NT; RSPO3-EGFP (RG)* alleles used for bulk transcriptome analysis. NT^+^RG^−^ (cortical cells, NT), NT^+^RG^+^ (CapLCs) and NT^−^RG^−^ (double negative, DN) cells were collected. SV20-8C12 NT iPSCs (parental line of SV20R2C4) were used as a negative control for RG signal detection. **(G)** Expression of key Cap markers in the indicated samples as described in (F). Mean + standard deviation. *p<0.05, **p<0.01, ***p<0.001. **(H)** Scatter plot comparing the average gene expression values between NT^+^RG^−^ cortical cells and NT^+^RG^+^ CapLCs. Differentially expressed genes (DEGs) (log-fold change >1, FDR <0.05, log2[RPKM+1] >1) are highlighted in color. Representative DEGs and their Gene Ontology (GO) enrichments are shown on the right. **(I)** Previously identified DEGs of human fetal adrenal Cap cells ^13,26^, compared to other fetal adrenal cell types were highlighted in red in the scatter plot shown in (H). **(J)** Heatmap showing upregulated DEGs of the indicated cell types identified from a multi-group comparison among CapLCs, NT, and DN (log-fold change >4, FDR<0.01). Enriched GO terms for DEGs are shown on the right. **(K)** Schematic of encapsulation of APLCs by CapLCs (left). The encapsulated APLCs were cultured without cytokines until mt21. BF and fluorescence images of the organoids at mt21 are shown on the right. Scale bar, 500 µm. **(L)** FACS plots of organoids at mt21 with or without encapsulation by CapLCs as shown in (K). **(M)** IF images of encapsulated organoids at mt21 showing RG, CD10 (green), NT (red), and DAPI (gray). Scale bar, 50 µm.

To identify cellular sources providing endogenous WNT signaling, we FACS-sorted NT^+^ APLCs at floating culture day (fl)18 (by removing NT^−^ non-cortical cells) and induced them into adrenal organoids. In contrast to organoids derived from unfractionated populations, organoids from NT^+^ cells failed to generate CD10^+^ DZLCs, suggesting that NT^−^ non-cortical cells provide essential WNT signaling for DZLC induction (Fig. 1E). Among non-cortical fractions in organoids, we previously identified RSPO3^+^ capsule (Cap)-like cells (CapLCs) bearing striking transcriptional similarities to Cap cells in human prenatal adrenal gland ^26^. CD10^+^ DZ cells first emerge within the parenchyma of the AP subjacent to these Cap cells at 6 wpf (fig. S1E, F). Thus, we hypothesized that RSPO3^+^ CapLCs within the organoid NT^−^ fraction serve as a source of WNT signaling required to specify DZLCs from APLCs. To visualize CapLCs in vitro, we established NT; RSPO3-EGFP (RG) dual fluorescence reporter NTRG iPSCs (designated as SV20R2C4) (fig. S1G, H). At Matrigel culture day (mt)21 following fl18 [fl18mt21], organoids induced from NTRG reporter iPSCs showed induction of NT^−^RG^+^ cells along with NT^+^ cortical cells and NT^−^RG^−^ cells (Fig. 1F, fig. S1I). Comparison of FACS-sorted cell fractions by bulk-RNA-seq revealed that RG^+^ cells upregulate markers of Cap cells previously identified in both human and mouse adrenal glands (e.g., *RSPO3*, *GLI1*, *GLI2*) (Fig. 1G-J, Table S1, 2). NT^−^RG^+^ cells differentially expressed genes (DEGs) that were enriched with GO terms such as “smoothened signaling pathway” or “canonical Wnt signaling pathway”, confirming their identity as CapLCs (Fig. 1H, J) ^15,19^. RSPO3 was uniquely expressed in CapLCs and most abundant among all RSPO family (fig. S1J). Notably, encapsulation of FACS-sorted NT^+^ APLCs by RG^+^ CapLCs resulted in robust induction of CD10^+^ DZLCs in vitro, suggesting that CapLCs drive canonical WNT signaling in APLCs, thereby inducing DZLCs (Fig. 1K-M).

### Reconstitution of the niche allows robust induction of DZLCs in vitro

Given the critical role of CapLC-derived RSPO3 in DZLC induction in our unfractionated organoids, we next assessed the sufficiency of RSPO3 in DZLC induction from purified NT^+^APLCs. As expected, FACS-sorted and reaggregated NT^+^ APLCs alone were insufficient to give rise to DZLCs (Fig.1L, 2A). However, further addition of RSPO3 induced DZLCs in dose-dependent manner (Fig. 2A). Reflecting shared domain organization, other members of RSPO family (i.e., RSPO1, RSPO2, RSPO4) were equally capable of inducing DZLCs (fig. S2A, B) ^27^. DZLC induction was further enhanced by addition of SB431542 (ACTIVIN/NODAL inhibitor), but suppressed by DAPT (NOTCH inhibitor), suggesting that such signaling might modulate DZLC induction (fig. S2C, D). In the presence of both RSPO3 and SB431542 (DZ medium), DZLC first appeared at mt7 and were progressively induced by mt21 (fig. 2E, F). The responsiveness of NT^+^ cells to inductive signaling first appeared at fl15, but was markedly increased by fl22 (fig. S2G). Histologically, DZLCs closely resembled DZ cells in vivo, present in tightly packed clusters and exhibiting scanty basophilic cytoplasm with round/ovoid nuclei (Fig. 2B). In contrast to FZ cells, ultrastructural analysis revealed that cytoplasm of DZ/DZLCs contain significantly fewer mitochondria and osmiophilic lipid granules (Fig. 2B). Moreover, DZLCs exhibited striking immunophenotypic and transcriptional similarities to DZ cells, with strong expression of MME, NOV and LEF1 and high MKI67 proliferation index, whereas they showed no/markedly diminished expression of key FZLC/steroidogenic markers (i.e., STAR, CYP11A1, CYP17A1, SULT2A1) (Fig. 2C, D, fig. S2H). As robust DZLC induction was observed in organoids derived from two additional NT reporter iPSC lines (NT 679Sev4-2123 [designated as 2123] and WGNT SV20-211[designated as 211], see METHODS) as well as non-reporter iPSCs (Penn123i-SV20, designated as SV20) (fig. S2I-K), we have identified a highly reproducible and tractable method recapitulating the DZ specification process, allowing for functional assessment of human DZ state.

**Fig. 2.**
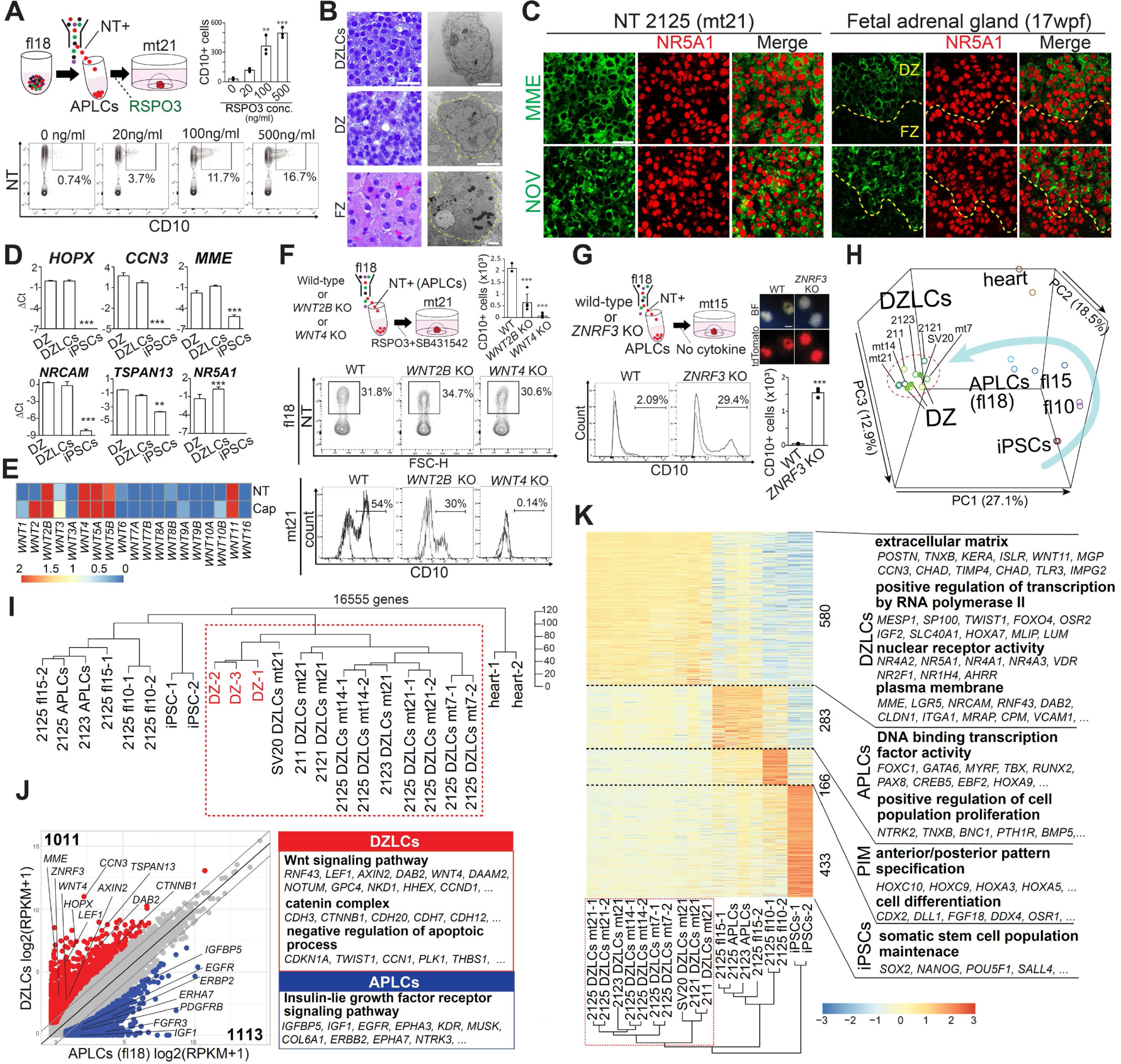
Reconstitution of the niche allows robust induction of DZLCs in vitro. **(A)** Schematic of DZLCs induction from APLCs (top left). FACS analysis showing the percentage (bottom) and total number per organoid (top right) of NT^+^CD10^+^ DZLCs at mt21 cultured with RSPO3 (0, 20, 100, 500 ng/ml). Cells were derived from 211 hiPSCs. Mean + standard deviation (n = 3). *p<0.05, **p<0.01, ***p<0.001 versus 0 ng/ml. **(B)** H&E images (left) and transmission electron microscope (TEM) images (right) of DZLCs at mt21, along with DZ and FZ cells from a fetal adrenal gland at 19 weeks post-fertilization (wpf). Scale bars, 20 μm (left); 1 μm (right). **(C)** IF images of DZLCs at mt21, DZ and FZ cells at 17 wpf, showing MME and NOV (green), NR5A1(red) with their merged images. Scale bar, 30 μm. **(D)** qPCR analysis of key genes in DZ (15 and 17 wpf), DZLCs and iPSCs. 2125 iPSCs were used. *p<0.05, **p<0.01, ***p<0.001 versus DZ. **(E)** Heatmap showing the expression of WNT ligands in CapLCs and NT^+^ cells at mt21, as shown in Fig. 1F. **(F)** FACS analysis of *WNT2B* (derived from 238A9) and *WNT4* mutant (derived from 237B11) organoids, along with the parental control (derived from 2125 iPSCs) at fl18 and mt21. The number of CD10^+^ cells at mt21 was assessed and data are presented as mean + standard deviation (n = 4). *p < 0.05, **p < 0.01, ***p < 0.001. **(G)** FACS analysis and images (BF and fluorescence) of *ZNRF3* mutant (derived from ZT1C3) organoids or parental control (derived from 2125 iPSCs) at mt15. Organoids were cultured without cytokines following FACS-sorting of NT^+^ APLCs at fl18. The percentage and total number of NT^+^CD10^+^ cells were assessed. Data are presented as mean ± standard deviation (n = 3). *p < 0.05, **p < 0.01, ***p < 0.001. Scale bar, 200 μm. (**H, I**) Principal component analysis (PCA) (H) and unsupervised hierarchical clustering (UHC) using complete linkage clustering (I) of bulk transcriptomes during DZLC induction. A cluster containing DZ and DZLCs are outlined by dotted red lines. DZLCs were derived from 2125 (at mt7, 14, 21), 211, 2121, 2123, and SV20 (at mt21), and APLCs (at fl18) were derived from 2125 and 2123 iPSCs. iPSCs, fl10, and fl15 organoids were derived from 2125 iPSCs. DZ (FACS-sorted as in fig. S5G) and whole heart cells were derived from fetuses (15, 17, 19 wpf). DZLCs were FACS-sorted by CD10 expression, and organoids at fl15, 18 were isolated by NT expression. **(J)** Scatter plot comparing the average gene expression values between DZLCs and APLCs (fl18) derived from 2125 hiPSCs. DEGs (log2-fold change >1, FDR <0.05, log2[RPKM+1] >1) are highlighted and representative DEGs and their GO enrichments are shown on the right. **(K)** Heatmap showing upregulated DEGs (log-fold change >4, FDR <0.01) identified from a multi-group comparison among DZLC (outlined in dotted red lines), APLC (fl15, fl18), PIM (fl10), and hiPSC clusters. Hierarchical clustering of the indicated samples was performed using complete linkage clustering. DEGs and their Gene Ontology (GO) enrichments are shown on the right.

RSPO3-mediated DZLC induction was significantly diminished when CRISPR/Cas9 targeting *CTNNB1* (encoding β-catenin, key mediator of canonical WNT signaling) were introduced by lentivirus vectors, further confirming its dependency on canonical WNT-signaling (fig. S3A). Because RSPO3 potentiates canonical WNT signaling through increased membrane availability of Frizzled (FZD) (i.e., receptors for WNT ligands), its action is considered to be dependent on WNT ligands ^7^. Thus, we next evaluated the requirement of endogenous WNT ligands in RSPO3-mediated DZLC induction. Organoids cultured with RSPO3 and SB431542 (DZLC medium) failed to generate DZLCs in the presence of LGK974 (an inhibitor for porcupine, a membrane-bound O-acyltransferase critical for secretion of WNT ligands), further suggesting the critical role of endogenous WNT ligands. Among WNT ligands activating canonical WNT signaling, NT^+^ cells expressed *WNT2B* and *WNT4*, previously identified as functional WNT ligands in mouse adrenal cortex (Fig. 2E) ^28,29^. Thus, we generated both *WNT2B* and *WNT4* homozygous mutant iPSC lines using CRISPR/Cas9 (fig. S3C-E). While both lines were readily induced into NT^+^ APLCs, we noted partial or complete loss of DZLC induction in *WNT2B* or *WNT4* mutant lines, respectively, suggesting dependency on autocrine/paracrine WNT2B and WNT4 in RSPO3-mediated DZLC induction (Fig. 2F).

RSPO3 binding to its receptors LGR4/5 promotes ubiquitin-mediated degradation of ZNRF3/RNF3, negative regulators of FZDs (i.e., WNT receptors), thereby increasing membrane availability of FZDs and potentiating WNT signaling. To assess the role of *ZNRF3* in RSPO3-mediated DZLC induction, we next generated bialleleic *ZNRF3* mutant iPSCs (fig. S3F, G). Strikingly, FACS-sorted NT^+^ APLCs from *ZNRF3* mutant iPSCs could be robustly induced into DZLCs even in the absence of RSPO3, and this effect was completely abolished in the presence of porcupine inhibitor, LGK974 (Fig. 2G, fig. S3F-H). Together, our finding strongly indicates that RSPO3-mediated DZLC induction is dependent on both ZNRF3 and endogenous WNT ligands.

### Transcriptional dynamics of DZLC induction and resemblance between DZLCs and DZ in vivo

To capture the transcriptional dynamics of the DZLC induction process, we conducted bulk RNA-seq on different stages of induction (iPSCs, fl10, fl15, fl18 [APLCs], DZLCs [fl18mt7, fl18mt14, fl18mt21]) using 2125 iPSCs along with DZLCs induced from various reporter and non-reporter iPSC lines (male: 211, 2123, SV20; female: 2121, 2125). We then compared them with FACS-sorted CD10^+^ DZ cells derived from human prenatal adrenal gland in vivo. PCA analysis demonstrated progressive differentiation of iPSCs into APLCs, then into DZLCs over time (Fig. 2H). Notably, DZLCs closely clustered with in vivo DZ cells in both PCA and unsupervised hierarchical clustering (UHC) analyses, highlighting striking transcriptional similarities (Fig. 2I). Concordantly, key DZ marker genes (e.g., *HOPX*, *NOV*, *MME, TSPAN13* as assessed by qPCR) were expressed at remarkably similar levels in DZ cells and DZLCs (Fig. 2D, fig. S3I). DEGs analysis revealed that DZLCs upregulated genes associated with WNT signaling (e.g., *LEF1*, *AXIN2*, *TCF4*), extracellular matrix (e.g., *POSTN*, *TNXB*) or nuclear receptors (e.g., *NR4A2*, *NR5A1*, *VDR*), suggesting that these genes are involved in maintaining their unique progenitor function (Fig. 2J, K, fig. S3J, Table S3, 4). Together, these findings suggest that iPSC-derived organoids progressively acquire a DZLC transcriptional state highly resembling that of DZ cells in vivo.

### Coordinated action of canonical WNT signaling and ACTH mediates transdifferentiation of human prenatal adrenal cortex

Histomorphological observations indicate that DZ cells may contribute to the maintenance of human prenatal adrenal cortex homeostasis ^4,30^. These cells are proposed to have a role in self-renewal and potentially undergo centripetal trans-differentiation into steroidogenic cells. To test this concept experimentally and to identify potential signaling principles driving trans-differentiation, we set out to compare the trans-differentiation potential of DZ cells and DZLCs. In prenatal adrenal cortex at the second trimester, CD10^−^SULT2A1^−^HSD3B2^+^ TZ cells were observed between CD10^+^ DZ cells and SULT2A1^+^ FZ cells, and were also positive for key steroidogenic proteins with variable intensity (i.e., STAR, CYP11A1, CYP11B1, CYP17A1) (Fig. 3A, B, fig. S4A-B), suggesting that the DZ transdifferentiates into the FZ through a TZ intermediate. As zG-to-zF trans-differentiation in mice is driven by ACTH ^21^, and the key WNT-target gene, *LEF1* in the DZ remains expressed in TZ cells in humans (Fig. 3C, fig. S4C), we hypothesized that a combination of ACTH and RSPO3 would promote trans-differentiation of DZLCs into TZLCs. In support of this notion, we found that DZLCs (both fresh and freeze-thawed) differentiate into TZLCs by upregulating HSD3B2 and CYP17A1 and producing high levels of glucocorticoids (i.e., cortisol) when treated with RSPO3 and ACTH (TZ medium) (Fig. 3D-H). Moreover, this process occurred through PKA signaling and dose-dependent action of ACTH (fig. S4D-F). Notably, transient stimulation of TZLCs with ACTH (after two days of culturing in ACTH-depleted medium) markedly enhanced cortisol production, confirming their ACTH responsiveness and functional maturation (Fig. 3I). Finally, we determined if this platform could be exploited to functionally interrogate human glucocorticoid biosynthetic pathways, which heretofore could only be assessed indirectly. As proof of principle, we used Trilostane, an inhibitor for HSD3B2, as well as Metyrapone and Osilodrostat, inhibitors of 11β-hydroxylase (i.e., CYP11B1, CYP11B2), both of which are required to generate cortisol from progestogens. All inhibitors significantly suppressed the production of cortisol, supporting the potential utility of this platform as a tool for screening inhibitors of glucocorticoid biosynthesis (Fig. 3J).

**Fig. 3.**
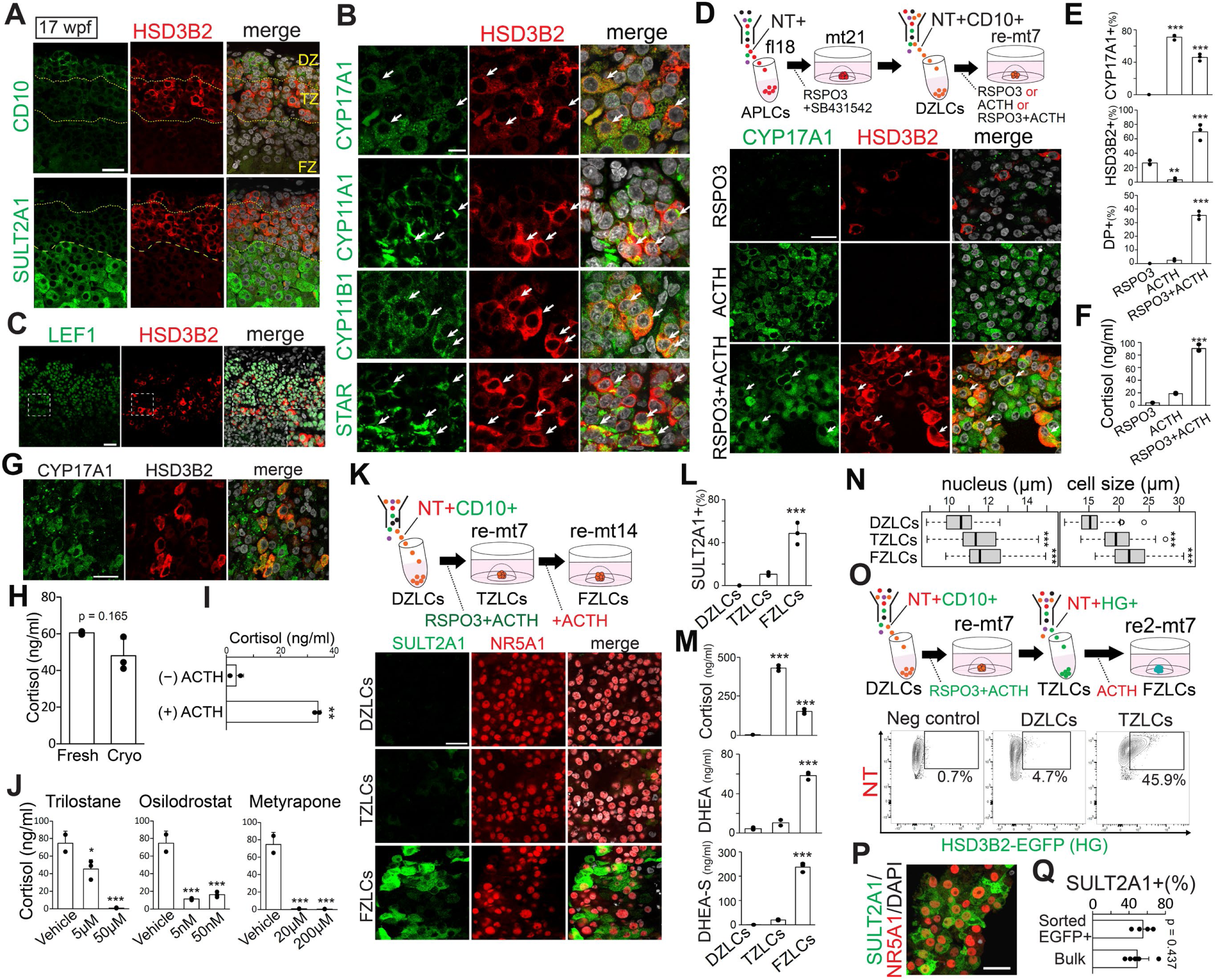
Coordinated action of canonical WNT signaling and ACTH mediates transdifferentiation of human prenatal adrenal cortex. **(A)** IF images of fetal adrenal glands at 17 wpf showing CD10, SULT2A1 (green), and HSD3B2 (red), along with their merged images with DAPI (gray). Scale bar, 30 μm. **(B)** High magnification IF images of the TZ at 17 wpf. Double-positive cells for the indicated markers are shown by arrows. Scale bar, 10 μm. **(C)** IF images of a fetal adrenal gland at 17 wpf for the indicated markers. High magnification merged images of the indicated region are shown as an inset, highlighting cells reactive for both LEF1 and HSD3B2. Scale bar, 50 μm. **(D)** IF images of organoids derived from 2125 iPSCs cultured with RSPO3 (100 ng/ml), ACTH (5 nM), or both for an additional seven days (re-mt7) following DZLC induction and FACS purification of NT^+^CD10^+^ DZLCs (mt21). CYP17A1 (green), HSD3B2 (red), and their merged images with DAPI are shown. **(E)** Quantification of IF analysis showing the percentages of CYP17A1^+^, HSD3B2^+^, and double positive (DP) cells among NR5A1^+^ cells, as described in (D). **(F)** Cortisol production as described in (D). Culture supernatant was collected at 72 h, and cortisol concentration was determined by LC-MS/MS. Data are presented as mean + standard deviation (n = 3). *p < 0.05, **p < 0.01, ***p < 0.001 versus RSPO3. **(G)** IF images of TZLCs induced from cryopreserved DZLCs after seven days of RSPO3 and ACTH treatment. CYP17A1 (green), HSD3B2(red), and their merged images are shown. Scale bar, 30 μm. **(H)** Cortisol production by TZLCs as described in (G). TZLCs induced from fresh (non-frozen) DZLCs were used as a control. Culture medium was collected at 72 h. Data are presented as mean + standard deviation (n = 3). **(I)** Cortisol production following transient ACTH treatment. TZLCs derived from 2125 iPSCs were cultured for one day after ACTH withdrawal, followed by stimulation with or without ACTH for an additional day. RSPO3 (100 ng/ml) was maintained in all conditions. Culture supernatants were collected at 24 h. Data are presented as mean + standard deviation (n = 2). *p < 0.05, **p < 0.01, ***p < 0.001. **(J)** Cortisol production in TZLCs treated with Trilostane, Osilodrostat, or Metyrapone. A vehicle control was treated with DMSO. Culture supernatants were collected at 72 h, and cortisol concentration was measured. Data are presented as mean + standard deviation (n = 3, except vehicle: n = 2). *p < 0.05, **p < 0.01, ***p < 0.001 versus vehicle control. **(K)** Trans-differentiation of DZLCs to FZLCs through TZLCs. FACS-sorted DZLCs were cultured with RSPO3 and ACTH for seven days (re-mt7, TZLCs), followed by ACTH only for an additional 7 days (re-mt14, FZLCs). Immunofluorescence (IF) images of indicated cells showing SULT2A1 (green), NR5A1 (red), and their merges with DAPI (gray). Scale bar, 30 μm. **(L)** Quantification of SULT2A1 positive cells in (K). Mean + standard deviation (n = 3). *p<0.05, *p<0.01, ***p<0.001 versus DZLCs. **(M)** Cortisol, DHEA, and DHEA-S production in (K). Mean + standard deviation (n = 3). *p<0.05, *p<0.01, ***p<0.001 versus DZLCs. **(N)** Boxplots showing IF quantification of nucleus and cell size (n = 100) of DZLCs, TZLCs, and FZLCs derived from 2125 iPSCs as described in (K). *p<0.05, **p<0.01,***p<0.001 versus DZLCs. **(O)** FACS plots of NT^+^HSD3B2-EGFP (HG)^+^ TZLCs at re-mt7 (middle). Cells were derived from MH2C29 (NTHG) and 2125 (NT, parental line of MH2C29, negative control). **(P)** Immunofluorescence (IF) of FACS-sorted NT^+^HG^+^ TZLCs treated with ACTH (5 nM) for seven days (re2-mt7) as described in (O). Scale bar, 30 μm. **(Q)** Quantification of SULT2A1 positive cells in organoids as described in (P). FZLCs induced from non-sorted bulk organoids are shown as positive control. Cells were derived from MH2C29 (NTHG) (n = 4).

Given that RSPO3 acts locally ^16^, the inner cortex distal to the capsule is likely spared from RSPO3 action, leading us to hypothesize that removal of RSPO3 would drive differentiation of TZLCs into functional FZLCs. In support of this idea, removal of RSPO3 and treatment of TZLCs with ACTH alone for an additional seven days enhanced the expression of FZ markers (e.g., SULT2A1, POR) accompanied by cellular hypertrophy and increased adrenal androgen production (i.e., DHEA, DHEA-S). Concomitantly, HSD3B2 expression was downregulated (Fig. 3K-N, fig. 4G-I). These findings suggest successful differentiation of TZLCs into functional FZLCs. Importantly, FACS-sorted HSD3B2-EGFP^+^ TZLCs could give rise to FZLCs upon reaggregation culture, further lending support that HSD3B2^+^ TZLCs, but not contaminating immature DZLCs, give rise to FZLCs (Fig. 3O-Q, fig. S4J, K). Notably, the trans-differentiation potential of DZLCs is maintained even after long-term culture of in DZ medium (fl18mt50), consistent with progenitor characteristics capable of long-term self-renewal (fig. S5A-F).

**Fig. 4.**
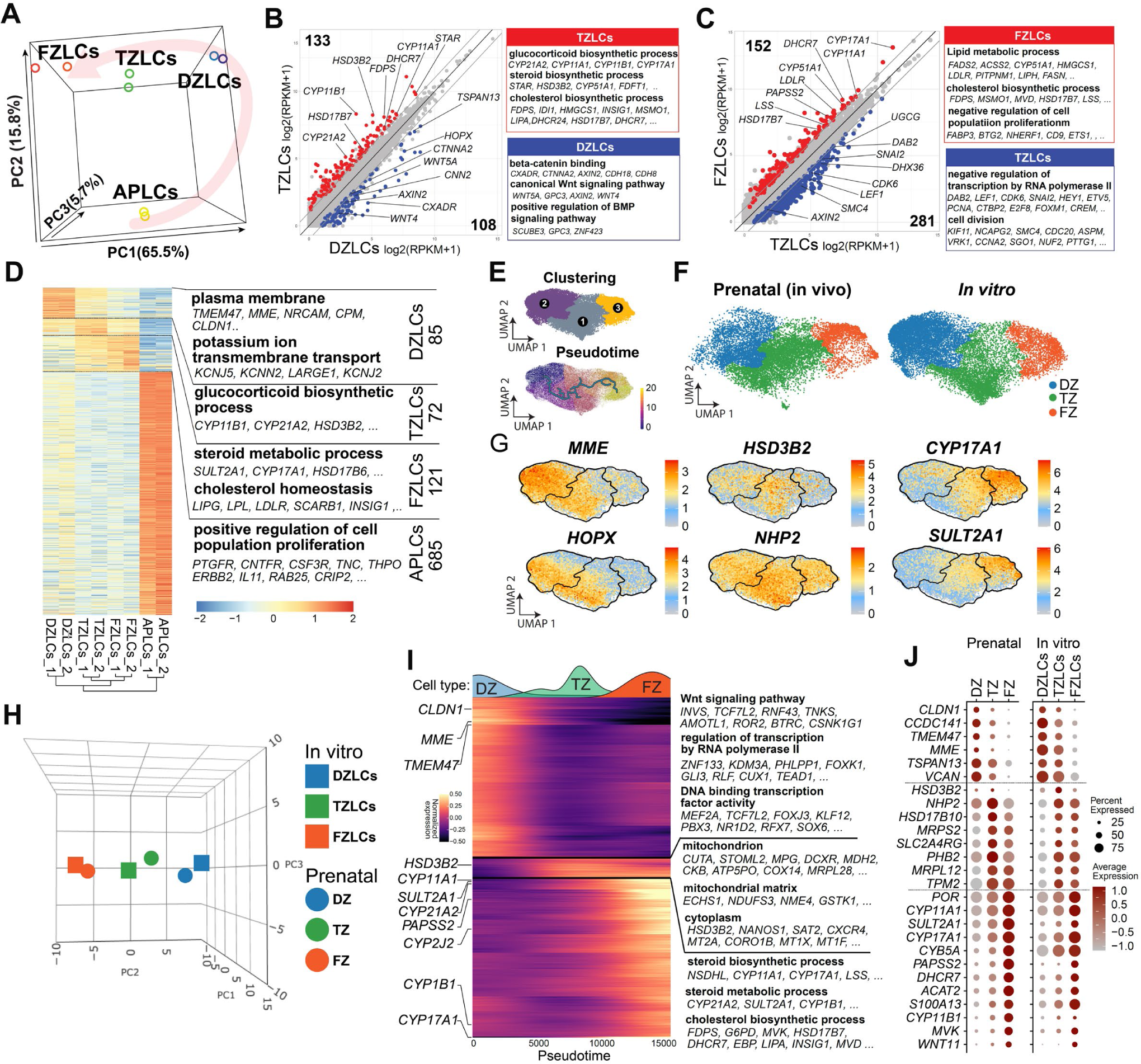
Transcriptional dynamics during human adrenocortical trans-differentiation. (**A**) PCA analysis of bulk transcriptomes from APLCs, DZLCs, TZLCs, and FZLCs derived from 2125 hiPSCs. (**B, C**) Scatter plots comparing the average gene expression between DZLCs and TZLCs (b) or TZLCs and FZLCs (C). DEGs (log-fold change >1, FDR <0.05, log2(RPKM+1) >1) are highlighted in colors. Representative genes and their GO enrichments are shown on the right. **(D)** Heatmap showing DEGs (log2-fold change >3, FDR<0.01) identified by a multi-group comparison among APLCs, DZLCs, TZLCs, and FZLCs. Enriched GO terms among representative DEGs are shown on the right. **(E)** UMAP plots of single-cell transcriptomes showing clusters (top) and pseudotime progression of integrated in vitro and prenatal adrenocortical cells (bottom). **(F)** Prenatal (left) and in vitro-derived (right) adrenocortical cells from UMAP plot in (E). Cells were colored according to clusters annotated by marker genes as in (G). **(G)** Expression of representative marker genes projected on the UMAP plot as in (E). **(H)** PCA analysis of average expression values (pseudo-bulk) of indicated cell types defined in (F). **(I)** Heatmap of gene expression along pseudotime trajectory as in (E). Top 200 genes selected based on percentage difference in expression, with a minimum percentage difference of 0.08, log2-fold change > 0.585 and q-value < 0.05. **(J)** Dot plot of key gene (DEG) expression by cell type. The size of the dot indicates the percentage expression of the gene in question for that cell type and color displays the mean log2 fold-change as compared with all other cell types.

Having determined conditions driving DZLC trans-differentiation in vitro, we next validated our finding using DZ cells in vivo. First, we FACS-sorted CD10^+^ DZ cells and CD10^−^ TZ/FZ cells as described previously (fig. S5G, H) ^26^. We found that FACS-sorted and reaggregated DZ cells maintain DZ markers ex vivo for two weeks in the presence of DZ medium, suggesting that, like DZLCs, DZ maintenance is dependent on RSPO3 (fig. S5I, J). Importantly, CD10^−^ TZ/FZ cells failed to upregulate DZ markers ex vivo even in the presence of DZ medium suggesting unidirectionality of trans-differentiation (fig. S5I, J). In addition, we found that DZ cells can readily differentiate into TZ-like cells ex vivo upon culturing in TZ medium (ACTH and RSPO3) and FZ-like cells by additional culture in FZ medium (ACTH alone), similar to DZLC trans-differentiation (fig. S5K-O). Altogether, these findings suggest that DZ/DZLCs induced and maintained by RSPO3, sequentially trans-differentiate into steroidogenic TZ and FZ cells through concerted action of RSPO3 and ACTH.

### Transcriptional dynamics during human adrenocortical transdifferentiation

To gain insights into lineage trajectory during DZ trans-differentiation, we performed bulk RNA-seq. PCA analysis revealed a directional trajectory in which DZLCs progress towards FZLCs through TZLCs along PC1 axis (Fig. 4A). Consistent with cortisol production by TZLCs, DZLC-to-TZLC progression was characterized by upregulation of genes related to glucocorticoid steroid synthesis (e.g., *CYP21A2*, *CYP11A1*, *HSD3B2*, *CYP17A1*), which were enriched in GO terms such as “glucocorticoid biosynthetic process” (Fig. 4B, Table S5). As TZLCs further progressed into FZLCs, there was further upregulation of genes involved in steroid biosynthesis that were enriched with GO terms such as “lipid metabolic process” or “cholesterol biosynthetic process” (Fig. 4C, D, Table S5). Moreover, FZLCs revealed high levels of expression of genes related to adrenal androgen biosynthesis such as *CYP17A1*, *SULT2A1* or *PAPSS2* consistent with active androgen production (Fig. 4D). Along the trajectory, key DZ markers (e.g., *HOPX*, *TSPAN13*) and WNT-target genes (e.g., *LEF1*, *AXIN2*) were gradually downregulated (Fig. 4B, C).

scRNA-seq data revealed a trajectory concordant with bulk RNA-seq where CD10^+^ DZLCs progressed into SULT2A1^+^ FZLCs though HSD3B2^+^ TZLCs (Fig. 4E-G). To assess whether adrenocortical trans-differentiation reconstituted in vitro follows what occurs in vivo, we integrated single cell transcriptomes from in vitro and in vivo prenatal adrenal samples. This analysis revealed that all samples intermingled with each other and shaped a continuous trajectory in which clusters were formed according to cell type (i.e., DZLCs/DZ, TZLCs/TZ and FZLCs/FZ) rather than sample origin (Fig. 4E, F). The transcriptional similarity was further confirmed by PCA analysis and marker gene expression, which was highly conserved between in vitro and in vivo cell types (Fig. 4H-J). Together, these findings reveal that adrenocortical trans-differentiation processes can be largely recapitulated in vitro using iPSCs.

### NR0B1 drives DZLC induction and safeguards against direct FZLC differentiation from APLCs

Having established a DZLC induction platform, we next assessed the transcription factors responsible for DZ induction. To this end, we found *NR0B1* and *GATA6* are among the most highly expressed transcription factors in both DZ and DZLCs (fig. S6A, Table S6). *NR0B1* mutation is associated with a form of primary adrenal insufficiency called X-linked adrenal hypoplasia congenita (AHC) ^31–33^. Most patients with AHC present with highly hypoplastic adrenal cortex consisting primarily of polygonal/cytomegalic cells resembling the FZ. As the function of *NR0B1* in human adrenal development and molecular pathophysiology of AHC remains largely unknown, we genetically assessed *NR0B1* function using our DZ induction system (fig. S6B-E). Although neither homozygotic (originated from 2125 iPSCs, XX) nor hemizygotic (originated from 211 iPSCs, XY) *NR0B1* mutation affected the induction of NT^+^ APLCs at fl18, mutant APLCs exhibited upregulation of genes related to steroid biosynthesis process, consistent with overall anti-steroidogenic role of *NR0B1* (Fig. 5A, fig. S6F, G) ^31^. A small subset of these genes was downregulated in our previously described *NR5A1* mutant APLCs (e.g., *CYP21A2*, *MC2R*, *MRAP*)^26^, supporting a potential antagonistic role of *NR0B1* in regulating key *NR5A1*-target genes (fig. S6H, Table S8).

**Fig. 5.**
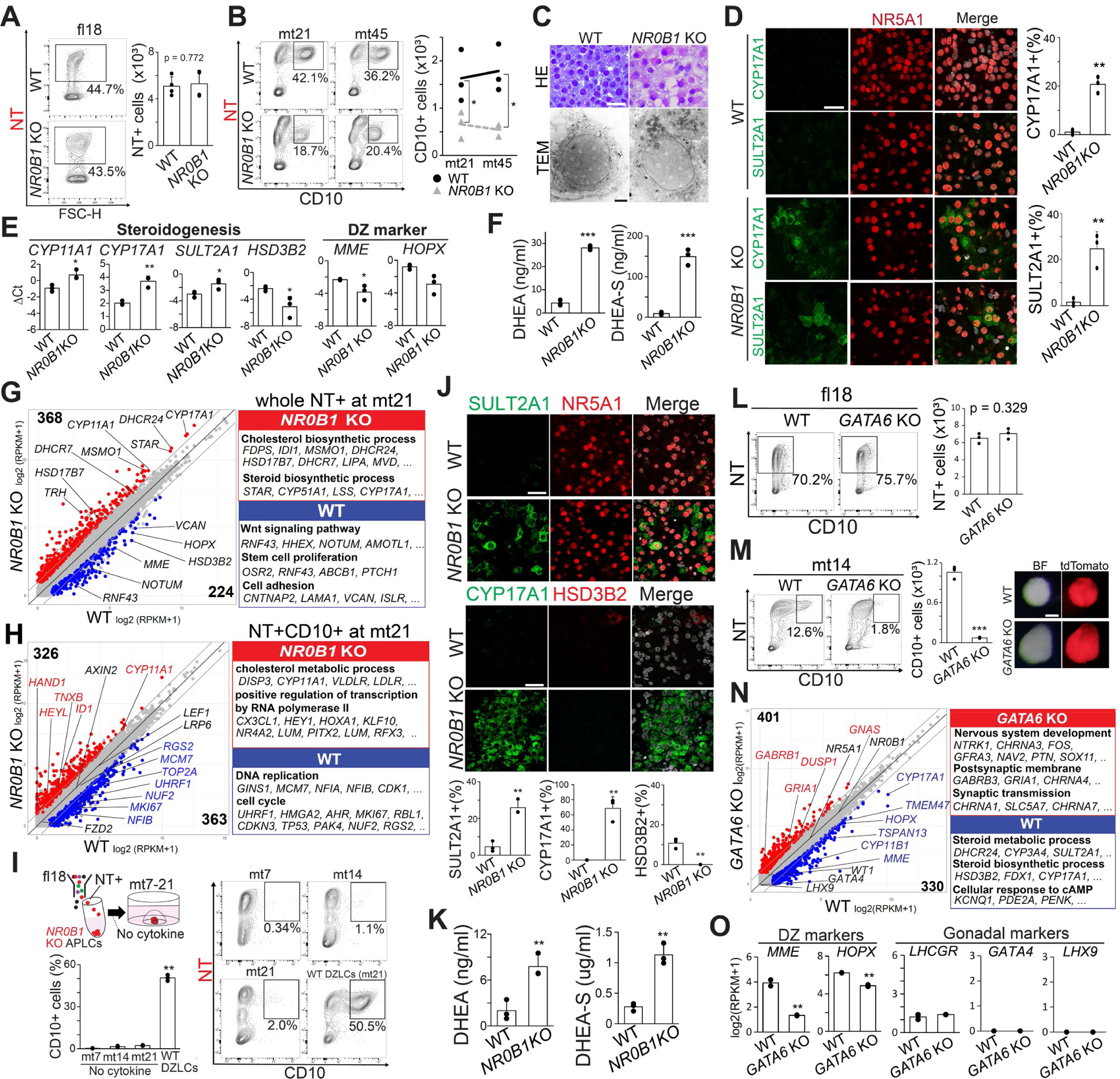
Critical role of *NR0B1* and *GATA6* in DZLC differentiation. **(A)** FACS plots of organoids at fl18 (APLCs), derived from 2125 (wild-type, WT) or 231C3 (*NR0B1* knockout, KO) hiPSCs. The total number of NTL cells per organoid was quantified. Data represent the mean + standard deviation (n = 4). **(B)** FACS plots of indicated organoids (left) and total number (right) of NTLCD10L cells per organoid at mt21 and mt45. Organoids were cultured in DZ medium following FACS sorting of NTL APLCs at fl18. Data represent the mean ± standard deviation (n = 3). *p < 0.05, **p < 0.01, *p < 0.001. **(C)** HE staining (top) and TEM images (bottom) of wild-type (WT) and *NR0B1* knockout (KO) organoids at mt21, as in (B). Scale bars: 10 µm (top), 2 µm (bottom). **(D)** IF images of indicated organoids at mt21, cultured in DZ medium, stained for CYP17A1 and SULT2A1 (green), NR5A1 (red), and merged with DAPI (gray) (left). Scale bar: 30 µm. Quantification of CYP17A1L and SULT2A1L cells is shown (right). Data represent the mean + standard deviation (n = 3). *p < 0.05, **p < 0.01, *p < 0.001. **(E)** qPCR analysis of whole organoids at mt21 for indicated genes as in (B). *p<0.05, **p<0.01, ***p<0.001. **(F)** DHEA and DHEA-S production by organoids at mt21. Culture supernatants were collected after 72 hours. Data represent the mean + standard deviation (n = 3). *p < 0.05, **p < 0.01, *p < 0.001. (**G, H**) Scatter plots comparing the average gene expression of WT and *NR0B1* KO organoids (FACS-sorted NT^+^ [top] or NT^+^CD10^+^ [bottom] cells) as in (B). DEGs (log2-fold change >1, FDR <0.05, log2[RPKM+1] >1) are highlighted in colors. Representative DEGs and their GO enrichments are shown on the right. **(I)** FACS analysis of *NR0B1* KO (231C3) organoids cultured without cytokines following FACS-sorting of NT^+^ APLCs at fl18. DZLCs at mt21, induced by DZ medium and derived from 2125 (WT) hiPSCs, served as a positive control. Mean + standard deviation (n = 3). *p<0.05, **p<0.01, ***p<0.001 versus mt7. **(J)** IF images of organoids cultured without cytokines at mt21, as described in (I), stained for the indicated markers. Quantification of marker expression is shown at the bottom. Data are presented as mean + standard deviation (n = 3). *p < 0.05, **p < 0.01, ***p < 0.001. Scale bar, 30 μm. **(K)** DHEA and DHEA-S production in culture supernatants, as described in (J). Culture medium was collected at 72 hours, and concentrations were measured. Data are presented as mean ± standard deviation (n = 3). *p < 0.05, **p < 0.01, ***p < 0.001. **(L)** FACS analysis of organoids at fl18 (APLCs) derived from 211 (WT) or 211e3c7 (*GATA6* KO) hiPSCs. The number of NT^+^ cells was quantified. Mean + standard deviation (n = 3). **(M)** FACS analysis of organoids at mt14 cultured with DZ medium following FACS-sorting of APLCs at fl18. Organoids were derived from 211 (WT) and 211e3c7 (*GATA6* KO) hiPSCs. The number of CD10^+^ cells was quantified. Data are presented as mean + standard deviation (n = 3). *p < 0.05, **p < 0.01, ***p < 0.001. BF and fluorescence (NT, red) images of the organoids at mt14 are shown on the right. Scale bar, 200 μm. **(N)** Scatter plot comparing the average gene expression of FACS-sorted NT^+^ cells in mt14 organoids between 211 (WT) and 211e3c7 (*GATA6* KO) lines. DEGs (log-fold change >1, FDR <0.05, log2(RPKM+1) >1) are highlighted in colors. Representative DEGs and their associated GO enrichments are shown on the right. **(O)** Expression of key DZ and gonadal markers in WT and *GATA6* KO organoids at mt14, assessed by bulk transcriptome analysis as described in (N). Data are presented as mean + standard deviation (n = 2). *p < 0.05, **p < 0.01, ***p < 0.001.

Remarkably, further differentiation of APLCs into DZLCs was markedly diminished in *NR0B1* mutant cells (Fig. 5B, fig. S6F). Histologically and ultrastructurally, we found that mutant cells have voluminous eosinophilic cytoplasm with abundant and pleomorphic mitochondria and lipid granules, thus resembling FZ cells. This was in stark contrast to wild-type DZLCs which had scant cytoplasm (Fig. 5C). Concordantly, in mutant lines, we found downregulation of DZ markers (e.g., *MME*, *HOPX*) and upregulation of FZ markers (e.g., *CYP17A1*, *SULT2A1*), which was accompanied by production of adrenal androgens suggesting that *NR0B1* is essential for DZ induction and for prevention of FZ differentiation (Fig. 5D-G, fig. S6I, J, Table S7). Although a small number of CD10^+^ cells were formed in mutant lines, transcriptome analysis of those cells in comparison to wild-type DZLCs exhibited overt abnormalities; mutant cells downregulated genes related to cell proliferation, a hallmark of the self-renewal properties of the DZ (Fig. 5H, Table S7) ^2,13^. Moreover, while many transcription factors (e.g., *HEY1*, *HOXA1*, *CDX2*, *NR4A3*) were aberrantly upregulated in mutant cells, WNT target genes (e.g., *LEF1*, *AXIN2, FZD2*) were not affected (Fig. 5H). Thus, defective DZ differentiation in *NR0B1* mutant cells is unlikely to be due to altered WNT signaling, but is more likely related to abnormal transcriptional programs. Finally, mutant CD10^+^ cells failed to transdifferentiate into TZLCs when cultured in TZ medium, indicating that *NR0B1* is crucial for induction of differentiation-competent DZLCs (fig. S6K).

Through scRNA-seq trajectory analysis of early prenatal adrenal cortex, we previously demonstrated that the AP gives rise to the FZ either directly (direct pathway) or indirectly through the DZ (indirect pathway/trans-differentiation) ^13^. The culture scheme we previously established to drive FZLCs from FACS-sorted APLCs did not yield overt CD10^+^ DZLCs throughout its time course, suggesting that FZLCs are induced through the direct pathway (fig. S6L, M) ^26^. Interestingly, prominent FZ differentiation occurred in *NR0B1* mutant organoids, even when cultured in the absence of DZ medium (base medium only) (Fig. 5I-K, fig. S6N, O). Because DZLC induction was negligible under such culture condition, mutant lines appeared to spontaneously differentiate into FZLCs directly from APLCs (i.e., via direct pathway) rather than through trans-differentiation from DZLCs (fig. S6P). Together, these findings demonstrate that *NR0B1* supports DZ induction and safeguards the AP against activating direct FZLC differentiation program.

### Critical role of *GATA6* in DZLC differentiation

Next, we assessed the role of *GATA6*, a transcription factor implicated in adrenal steroidogenic function and cortical development in mice (fig. S6Q, R) ^34,35^. Although *GATA6* mutation did not affect induction of NT^+^ cells in APLCs, subsequent induction into DZLCs was severely compromised, with a complete loss of CD10^+^ DZLCs (Fig. 5L, M). Consistently, bulk RNA-seq of unfractionated mutant organoids revealed downregulation of DZ markers (e.g., *MME*, *HOPX*, *TSPAN13*) (Fig. 5N). In contrast to previous report in mutant mice, mutant organoids did not differentiate into the gonadal lineage (Fig. 5N, O) ^34^. However, mutant organoids did upregulate many genes related to neuronal development (enriched with GO terms such as “Nervous system development” or “synaptic transmission”), suggesting that GATA6 reinforces DZ differentiation in part through safeguarding against aberrant activation of neuronal programs (Fig. 5N). Because *GATA6* mutant organoids showed no alteration in either *NR5A1* or *NR0B1* expression, the observed phenotypes were not dependent on these core adrenal transcription factors (Fig. 5N). Taken together, we have identified NR0B1 and GATA6 as key transcription factors driving proper DZ fate determination.

### Establishment of homeostatic functional zonation through trans-differentiation of encapsulated human adrenal organoids in vivo

We next tested whether DZLCs can sustain functional adrenocortical homeostasis in vivo. We first FACS-sorted CD10^+^ DZLCs, which were then reaggregated and transplanted under the kidney capsule of immunodeficient NCG mice. Under such conditions, grafts gradually lost NT^+^ cells by one month of transplantation (fig. S7A). We hypothesized that this is due to a lack of appropriate niche signals provided by Cap cells ^16^. Therefore, we next encapsulated DZLCs or APLCs by FACS-sorted RG^+^ CapLCs using transient floating co-cultures. APLCs were included based on our previous observation that APLCs can differentiate into DZLCs by encapsulation with CapLCs in vitro (Fig. 1L, M). Remarkably, CapLC encapsulation of either DZLCs or APLCs significantly improved graft survival. Grafts consisted of large number of NT^+^ cortical cells of human origin (human mitochondria antigen [hMito]^+^), EGFP^+^ CapLCs and a meshwork of mouse-derived CD31^+^ blood vessels (hMito^−^) (Fig. 6A, B, fig. S7B-D). Accordingly, flow cytometry of the grafts at 2-3 months of transplantation confirmed the persistence of RG^+^ CapLCs and NT^+^ cortical cells including CD10^+^ DZLCs (Fig. 6C).

**Fig. 6.**
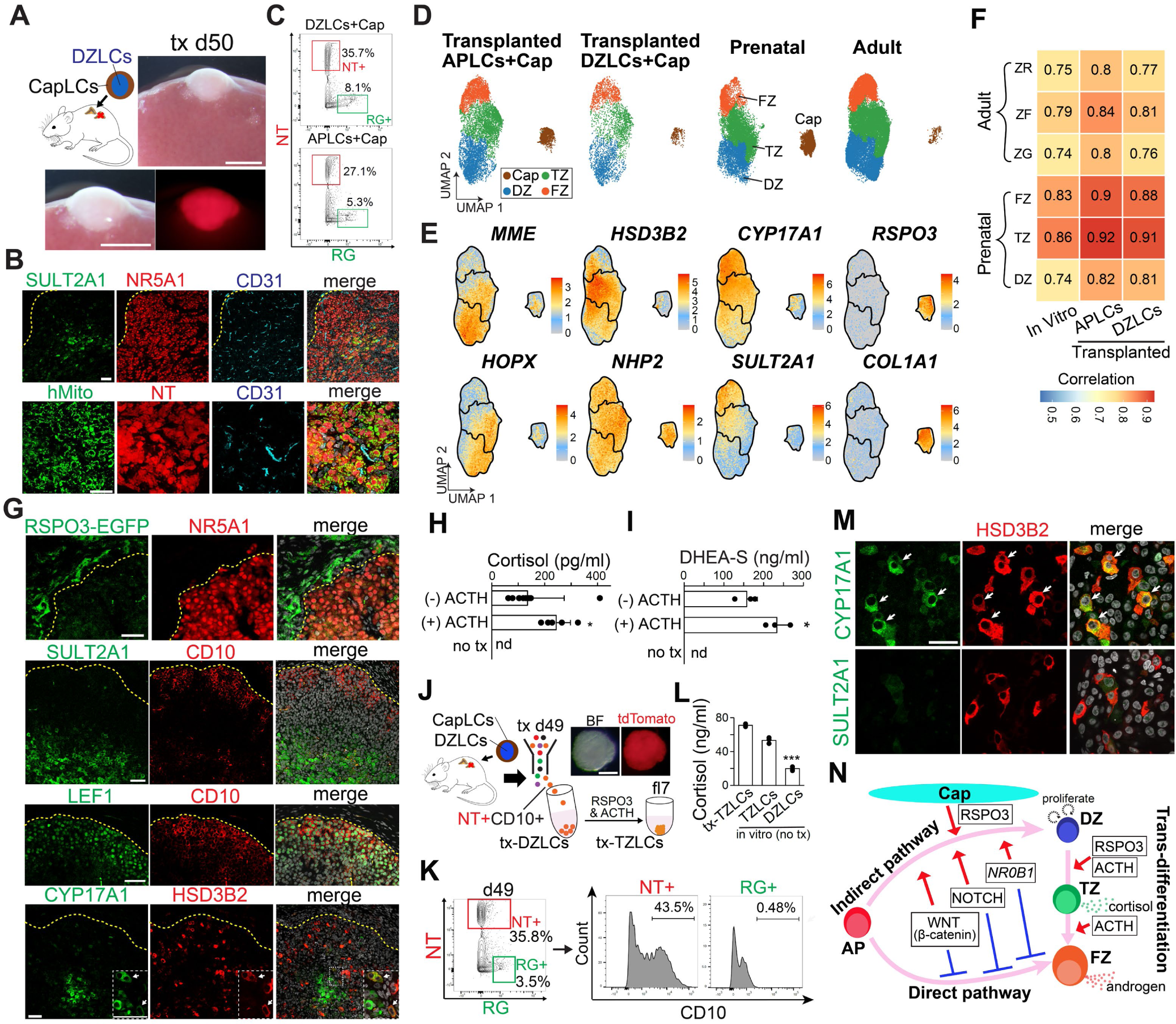
Establishment of homeostatic functional zonation through trans-differentiation of encapsulated human adrenal organoids in vivo. **(A)** CapLC-encapsulated DZLCs were transplanted under the kidney capsules of NCG mice that underwent hemi-adrenalectomy. DZLCs were derived from 2125 hiPSCs and CapLCs were derived from SV20R2C4 hiPSCs. Low-magnification BF image (top right) and high-magnification BF and fluorescence (NT, red) images (bottom) of organoids at d50 post-transplantation (tx d50). Scale bar, 2 mm. **(B)** IF images of encapsulated DZLCs at d50 post-transplantation, stained for indicated markers. Merged images with DAPI (gray) are shown on the right. hMito, human mitochondria antigen. Scale bar, 50 µm. **(C)** FACS plots of encapsulated DZLCs at d87 and APLCs at d90 post-transplantation. NT^+^ and RG^+^ cells were sorted for single-cell RNA-seq analysis. **(D)** UMAP plots showing sub-clustering of all cortical and Cap cells from the indicated samples. Cortical and Cap cells were isolated as described in fig. S7E, F, and then re-clustered. Cells are colored according to clusters annotated based on key markers, as shown in (E). **(E)** Expression of representative marker genes for the DZ (*MME*, *HOPX*), TZ (*HSD3B2*, *NHP2*), FZ (*CYP17A1*, *SULT2A1*) and Cap (*RSPO3*, *COL1A1*) projected on the UMAP plot in (D). **(F)** Pearson correlation of average expression values of transcriptomes. Cell clusters (i.e., DZ, TZ, FZ) of the indicated iPSC-derived samples were compared with their corresponding in vivo cell clusters on the left. Clusters were defined in (D). **(G)** IF images of encapsulated DZLCs at d50 post-transplantation, stained for the indicated markers. Merged images with DAPI (gray) are shown on the right. Yellow dotted lines denote the cortico-capsular border. Insets show magnified images of areas outlined by the white dotted line, highlighting CYP17A1 and HSD3B2 double-positive cells. Scale bar, 50 µm. **(H)** Serum cortisol levels following the cosyntropin stimulation test in mice that received encapsulated DZLCs. Transplanted mice were administered 100 µg/kg cosyntropin (ACTH) or PBS (no ACTH) 1 hour prior to blood collection. Non-transplanted control mice (no tx) received PBS and served as the negative control. Serum cortisol levels were measured by LC-MS. Data are presented as mean + standard deviation (n = 10 for no ACTH, n = 5 for ACTH, and n = 2 for control). *p < 0.05, **p < 0.01, *p < 0.001. nd, not detected. **(I)** Serum DHEA-S levels following the cosyntropin stimulation test in mice that received encapsulated DZLCs. Transplanted mice and control mice (no tx) were administered 100 µg/kg cosyntropin. Blood was collected before [(-) ACTH] and 1 hour after [(+) ACTH] cosyntropin injection. Data are presented as mean + standard deviation (n = 3). *p < 0.05, **p < 0.01, *p < 0.001. **(J)** Schematic illustrating the induction of TZLCs (tx-TZLCs) from FACS-sorted CD10^+^ DZLCs recovered from grafts derived from encapsulated DZLCs at d49 post-transplantation (tx-DZLCs). BF and fluorescence (NT, red) images of the organoids at fl7 are shown. Scale bar, 200 μm. **(K)** FACS analysis of NT^+^ or RG^+^ cells in grafts in (J) for their CD10 expression. **(L)** Cortisol production from TZLCs at re-fl15 induced in (j) (n = 3). *p<0.05, **p<0.01,***p<0.001. **(M)** IF images of tx-TZLCs at re-fl15 for CYP17A1, SULT2A1 (green), HSD3B2 (red), and their merged images with DAPI (gray). White arrows indicate double positive cells. Scale bar, 30 μm. **(N)** A model showing DZLCs induction and its trans-differentiation.

To further evaluate the differentiation potential of CapLC-encapsulated xenografts (APLCs or DZLCs), we next sought to elucidate their lineage trajectory and gene expression dynamics. To this end, we conducted scRNA-seq on FACS-sorted NT^+^ or RG^+^ cortical cells in xenografts after filtering out contaminated mouse cells (fig. S7E, Methods) ^36^. Clustering and projection of these cells into UMAP revealed two clusters, which could be readily identified as RSPO3^+^ CapLCs and NR5A1^+^ cortical cells (fig. S7F, G). A comparison of the transcriptomes of xenograft-derived cortical cells with in vivo human adrenal cortex revealed a remarkable similarity to both prenatal and adult adrenal cortical cells (Fig. 6D, E, fig. S7H, I). These findings suggest that transcriptional programs governing trans-differentiation is largely conserved between prenatal and adult cortex, and that both the DZLCs and APLCs are preprogrammed to establish homeostatic trans-differentiation. Notably, comparison of the transcriptome of xenograft-derived CapLCs clusters with both prenatal and adult Cap cells in vivo (Fig. 6D) suggested that CapLC identity was maintained after transplantation in vivo. However, in a cell type-by-cell type comparison, xenografted cells exhibited an even higher correlation to their in vivo counterparts (both prenatal and adult) than did cells trans-differentiated in vitro, suggesting that additional local and/or endocrine cues provided by in vivo environment play an important role in normal trans-differentiation (Fig. 6F).

In support of our transcriptional studies, immunofluorescence analysis revealed that grafts consisted of CD10^+^ DZLCs, CD10^−^HSD3B2^+^CYP17A1^+/−^ TZLCs and SULT2A1^+^CYP17A1^+^ FZLCs. Although zonation was not as distinct as native human adrenal cortex, these cell types were arranged in centripetal orientation (Fig. 6G, fig. S7J). Consistent with their trans-differentiation in vivo, we could detect cortisol and DHEA-S production in the serum of recipient mice (Fig. 6H, I, fig. S7K). Not surprisingly, these steroids were undetectable in non-transplanted mice because mouse adrenal glands do not possess significant CYP17A1 activity ^25,37^. Importantly, subcutaneous ACTH administration significantly increased both cortisol and androgen production suggesting that grafts retain ACTH responsiveness in vivo, a hallmark feature of the adrenal cortex (Fig. 6H, I). Finally, FACS-sorted CD10^+^ DZLCs recovered from the graft at three months of transplantation could be reaggregated and readily differentiated into cortisol-producing TZLCs in the presence of TZ medium (RSPO3 and ACTH), lending support that encapsulated DZLCs retain their differentiation potential even after long-term transplantation (Fig. 6J-M). Altogether, these findings support robust trans-differentiation potential of DZLCs enabling reconstitution of functional human adrenal cortex both in vitro and in vivo.

## Discussion

Here, we used iPSC-derived adrenocortical reconstitution and ex vivo culture of prenatal human adrenal cortex to characterize the cellular ontogeny of the human adrenal cortex and identify the underlying signaling and genetic mechanisms regulating its development. We provided evidence that two distinct pathways are operative in driving androgen-producing FZ in prenatal human adrenal cortex: 1) an indirect pathway in which the FZ originates from trans-differentiation of the DZ through a TZ intermediate and 2) direct pathway in which the FZ is induced directly from the AP (Fig. 6N).

NR0B1 is a nuclear receptor located on the X chromosome previously identified as a causative gene for AHC, a disease with early onset adrenal insufficiency ^1,31,32^. Biochemical analyses using immortal cell lines indicated its inhibitory role of steroidogenesis, particularly that mediated by NR5A1, a pro-steroidogenic master transcriptional regulator ^31,38^. Accordingly, *NR0B1* mutant mice exhibit early onset hyperactivation of adrenal steroidogenesis ^39,40^. However, in contrast to AHC patients, adrenal cortex development in these mice is normal. Consistent with the observed anti-steroidogenic function of NR0B1, we found that *NR0B1* mutant APLCs/DZLCs upregulate genes associated with steroidogenesis. Interestingly, most genes upregulated in mutant APLCs did not overlap genes downregulated as a consequence of *NR5A1* KO ^26^, suggesting that NR0B1 can regulate gene expression independent from NR5A1. Somewhat surprisingly, we discovered that NR0B1 regulates the dichotomous fate determination of the AP: by supporting DZ differentiation and suppressing direct FZ differentiation. Accordingly, *NR0B1* mutant APLCs exhibited exaggerated differentiation into FZLCs while failing to differentiate into functional DZLCs, consistent with the phenotype of AHC (i.e., neonatal-onset salt-wasting primary adrenal insufficiency). Notably, *NR0B1* mutant APLCs phenotypically resemble our previously described induction scheme in which APLCs are preferentially induced into FZLCs via the direct pathway (with minimal induction of DZLCs) by culture medium containing inhibitors of FGF, NOTCH and WNT signaling. In this study, we also demonstrated that inhibitors for NOTCH and WNT suppress DZ specification from APLCs. Thus, it is tempting to speculate that these signaling pathways might normally converge on NR0B1 to regulate dichotomous fate determination of the AP (Fig. 6N). This hypothesis warrants further investigation through biochemical and epigenetic approaches.

We demonstrated that DZLCs can be maintained long-term in the presence of a Cap-derived niche (i.e., RSPO3) while retaining their differentiation potential in vitro. The progenitor potential of DZLCs is further supported by ectopic xenotransplantation of CapLC-encapsulated DZLCs into mice where they maintain functional CD10^+^ DZLCs primarily at the periphery of the organoids while permitting their transdifferentiation into TZLCs and FZLCs. Importantly, without CapLCs, DZLCs were lost, and trans-differentiation did not occur, highlighting the critical role of Cap-derived niche factors for both progenitor maintenance and trans-differentiation.

Our in vivo organoids, capable of trans-differentiation, closely resemble both prenatal and adult adrenal cortex at the transcriptional level, indicating that the genetic program governing trans-differentiation is largely conserved across developmental stages and can be reconstituted using human iPSC-derived organoids. Remarkably, transplanted organoids survive long-term in ectopic locations and produce cortisol in response to ACTH, highlighting their potential as a cell-based therapy for primary adrenal insufficiency. Future preclinical studies in animal models will be crucial in validating this transformative approach.

## STAR Methods

### RESOURCE AVAILABILITY

#### Lead contact

Further information and requests for resources and reagents should be addressed to, and will be fulfilled by, the lead contact, Kotaro Sasaki.

#### Materials availability

hiPSCs generated in this study are available from the Lead Contact with a completed Materials Transfer Agreement.

#### Data and code availability

All data are available in the manuscript or the supplementary materials. scRNA-seq data have been deposited in the Gene Expression Omnibus (GEO) database (GSE291218).

### Experimental model and subject details

#### Collection of human adrenal glands

Fetal adrenal glands (5, 6, 8, 15, 17, and 18–19 weeks post-fertilization [wpf]) and hearts (9 wpf) used for bulk RNA-seq, single-cell RNA-seq, or histologic analyses were obtained from donors undergoing elective abortion at the University of Pennsylvania or Hokkaido University ^13^. All experimental procedures were approved by the Institutional Review Boards of the University of Pennsylvania (#832470) and Hokkaido University (#19-066). Informed consent being obtained from all donors for the use of aborted fetuses in research. Adult adrenal glands (ages 17, 38, and 50 years) used for single-cell RNA-seq were obtained from deceased donors through the International Institute for the Advancement of Medicine (IIAM), with informed consent obtained from the organ donor and/or their family for the use of adrenal glands in research.

Embryonic age was determined via ultrasonographic measurement of crown-rump length or head circumference. Fetal sex was identified using sex-specific PCR targeting the ZFX/ZFY loci. Fetal adrenal glands and hearts were submerged in RPMI-1640 medium and carefully dissected under a stereomicroscope to remove surrounding connective and adipose tissues.

#### Feeder free culture of human iPSCs

Parental hiPSC lines—Penn123i-SV20 (male), obtained from the iPSC Core Facility at the University of Pennsylvania, and Penn083i-679Sev4, derived from peripheral blood erythroblasts of a male donor, a kind gift from Dr. Wenli Yang—were used in this study ^41^. The 1390G3 line (female) was obtained from Dr. Masato Nakagawa at the Center for iPS Cell Research and Application, Kyoto University ^26,42^.

hiPSCs were maintained as previously described ^26^. Briefly, cells were cultured on 6-well plates (Thermo Fisher Scientific) coated with iMatrix-511 Silk (Nacalai USA) in StemFit Basic04 medium (Ajinomoto) supplemented with 50 ng/mL basic FGF (Peprotech) or StemFit Basic04CT (complete type) medium at 37 °C under 5% COL. For passaging or differentiation into adrenocortical lineages, hiPSCs at day 6–7 post-passaging were treated with a 1:1 mixture of TrypLE Select (Life Technologies) and 0.5 mM EDTA/PBS for 12–15 min at 37 °C to dissociate them into single cells. 10 µM ROCK inhibitor (Y-27632; Tocris) was added to the culture medium 24 hours post-passaging to enhance cell survival.

#### Authentication of cell lines used

All hiPSC lines used in experiments underwent G-banding analysis through Cell Line Genetics (Madison, WI). For genetically modified cells, master and secondary cell banks were established. First, cell lines were expanded until at least five cryovials were stored as a master cell bank. Next, a vial from the master bank was thawed and expanded to generate at least 20 cryovials as a secondary cell bank. All differentiation experiments were conducted using the secondary bank.

To ensure cell integrity, mycoplasma testing was performed using the MycoAlert detection assay (Cambrex). Passage-matched iPSCs (p35–p45) were used across all experiments. Cells were not passaged more than six times after thawing to minimize culture-induced genetic changes.

## Method details

### Generation of hiPSC lines bearing knock-in fluorescent reporter alleles

For donor vector construction to generate *NR5A1-3xflag-p2A-tdTomato* (*flag-NT*) alleles, homology arms spanning the 3’ end of the *NR5A1* locus (left arm: 1106 bp; right arm: 1328 bp) were PCR-amplified from genomic DNA of male hiPSCs (Penn123i-SV20) ^26,41^, and subcloned into the pCR2.1 vector using the TOPO TA Cloning Kit (Life Technologies). The *3xflag-p2A-tdTomato* fragment, along with the *Pgk-neo* cassette flanked by *loxP* sites, was PCR-amplified and inserted in-frame at the 3’-end of the *NR5A1* coding sequence using the GeneArt Seamless Cloning & Assembly Kit (Life Technologies). The *NR5A1* stop codon was removed to enable in-frame *p2A-NR5A1* protein expression.

Similarly, for donor vector construction to generate *RSPO3-p2A-EGFP* (*RG*) alleles, homology arms spanning the 3’ end of the *RSPO3* locus were PCR-amplified from genomic DNA of male hiPSCs (SV20-8C12) and subcloned into the *pCR2.1* vector. The *p2A-EGFP* fragment, containing a *Pgk-puro* cassette flanked by a *loxP* site, was PCR-amplified and inserted in-frame at the 3’-end of the *RSPO3* coding region.

For donor vector construction to generate *HSD3B2-p2A-EGFP* (*HG*) alleles, homology arms spanning the 3’ end of the *HSD3B2* locus were PCR-amplified from genomic DNA of male hiPSCs (NT 1390G3-2125) ^26^, and subcloned into the pCR2.1 vector. The *p2A-EGFP* fragment, containing a *Pgk-puro* cassette flanked by a *loxP* site, was PCR-amplified and inserted in-frame at the 3’-end of the *HSD3B2* coding region.

Pairs of single-guide RNAs (sgRNAs) targeting the 3’ end of *NR5A1*, *RSPO3*, and *HSD3B2* were designed using the Molecular Biology CRISPR Design Tool (Benchling). The sgRNA sequences were as follows: *NR5A1* 3’ end (5’-CTTGCAGCATTTCGATGAGC-3’ and 5’-CAGACTTGAGCCTGGGCCG-3’); *RSPO3* 3’ end (5’- GCTATCTCAACCAGATGCCC-3’ and 5’- ATGAGTCTACAATAATCTCA-3’); *HSD3B2* 3’ end (5’- GGCAACAGGGGCACAAGCCC-3’ and 5’- CCTTCTGTATGAAGCCAGGT-3’).

The sgRNAs were cloned into the *pX335-U6-Chimeric BB-CBh-hSpCas9n* (D10A) expression vector to generate the sgRNAs/Cas9n vector (a gift from Dr. Feng Zhang, Addgene #42335; http://n2t.net/addgene:42335; RRID: Addgene_42335) ^43^

To generate flagNT SV20-8C12 and NT 679Sev4-2123 hiPSCs, donor vectors (5 µg) and sgRNAs/Cas9n vectors (1 µg each) targeting the *NR5A1* locus were electroporated into one million hiPSCs (Penn123i-SV20, male and Penn083i-679Sev4, male) using a NEPA21 Type II Electroporator (Nepagene). Following neomycin selection, cells were transfected with a Cre recombinase-expressing plasmid (5 µg) to excise the *PGK-Neo* cassettes.

Cells were plated in 96-well plates (1 cell/well) using FACSAria Fusion single-cell plating mode. After 10–12 days of culture in StemFit Basic04 complete medium in 96-well plates and an additional 4–6 days in 12-well plates, cells were harvested. Half of the cells were used for genotyping, while the other half were cryopreserved.

To generate RGflagNT SV20R2C4 and HGNT MH2C29 hiPSCs, donor and sgRNAs/Cas9n vectors targeting *RSPO3* or *HSD3B2* loci were introduced into hiPSCs (flagNT SV20-8C12 or NT 1390G3-2125), respectively, followed by puromycin selection. Cells were then transfected with a Cre recombinase-expressing plasmid, subjected to single-cell plating via FACSAria Fusion, and screened for successful gene editing.

Targeted alleles, random integrations, or Cre-mediated recombination were identified via genotyping PCR using primers listed in Table S9. Sanger sequencing confirmed that the targeted alleles and non-targeted alleles were free of indels or unexpected mutations. Based on this screening, we found the following: flagNT SV20-8C12 hiPSCs were heterozygous for flagNT; NT 679Sev4-2123 hiPSCs were heterozygous for NT; RGflagNT SV20R2C4 hiPSCs were homozygous for RG; HGNT MH2C29 hiPSCs were heterozygous for HG. Some experiments used previously established hiPSC lines without modification including WG (WT1-p2A-EGFP);NT SV20-211 (WT1-p2A-EGFP, NT), NT 312-2121, and NT 1390G3-2125 ^26^.

### Generation of *RSPO3*, *WNT2B*, *WNT4*, *ZNRF3*, *NR0B1*, and *GATA6* knockout hiPSC lines

sgRNAs flanking the targeted exons were designed using the Benchling CRISPR design tool and cloned into the *pX335-U6-Chimeric BB-CBh-hSpCas9n* (D10A) expression vector to create the sgRNAs/Cas9n construct. The following sgRNA pairs were used: *RSPO3* (5’-TGATTTGATGGAGTTGGACA-3’ and 5’-GCAACATGCTCAGATTACAA-3’ for pre-exon2, 5’-AGACTATTTTTTGCTCTGGA-3’ and 5’-TGTCTCTCTTCATGTCCAAG-3’ for post-exon2); *WNT2B* (5’- ATATGCCATCTCATCAGCAG-3’ and 5’-AGAGGCAGCTTTTGTATATG for exon 3, 5’- ACTTCACCGCAGCCCGCCAA-3’ and 5’-AGAGGCAGCTTTTGTATATG-3’ for exon4); *WNT4* (5’-GGCCAAGCTGTCGTCGGTGG-3’ and 5’-GAGACGTGCGAGAAACTCAA-3’ for pre-exon2, 5’-CACTCGACTCCTTGCCCGTC-3’ and 5’-CTTGCCCGTCTTCGGCAAGG-3’ for post-exon2); *ZNRF3* (5’-ACTGGGCCTATGTAATAACA-3’ and 5’-GGACTTGTATGAATATGGCT-3’ for pre-exon2, 5’-GAATTGGACCCGAAACCATG-3’ and 5’-AAACCATGCCTCACTGTCCT-3’ for post-exon2); *NR0B1* (5’-GAGCGCGAAGCAAACGCGCG-3’ and 5’-CGCTCATAAGCATGTTGTAG-3’ for pre-exon1, 5’-CACGAGCACAAATCAAGCGC-3’ and 5’-TGCTCGTGGGCACGCAGTAG-3’ for post-exon1); *GATA6* (5’-CCATCCAGACGCCGCTGTGG-3’ and 5’-AGCCGCAGTTCACGCACTCG-3’ for exon 3).

Four pairs of sgRNAs/nCas9 vectors (2 μg each), except those targeting *GATA6* (two pairs), were electroporated into 1 million flagNT SV20-8C12 or NT 2125 hiPSCs (232B6, 238A9, 237B11, ZT1C3) using a NEPA21 Type II electroporator. After 48 hours, mCherry-positive cells were sorted as single cells into 96-well plates using a FACSAria Fusion flow cytometer. Clones were expanded, genotyped via PCR, and validated for indel formation through Sanger sequencing.

### Induction of DZLCs

The floating culture method for inducing APLCs followed a previously established protocol. DB27 medium (DMEM/F12 HEPES, 2% B27 supplement, 1% Glutamax, 1% insulin-transferrin-selenium [ITS-G], 1% MEM non-essential amino acids solution, 90 µM 2-mercaptoethanol, and 50 U/mL penicillin-streptomycin, all from Thermo Fisher Scientific) was used for fl0, fl1, and fl3. MK10 medium (Minimum Essential Medium alpha, 10% KSR, 55 µM 2-mercaptoethanol, and 100 U/mL penicillin/streptomycin, all from Thermo Fisher Scientific) was used for fl5, fl7, fl10, fl12, and fl15 ^26^.

Cytokine supplementation was as follows: fl0 medium includes 2 ng/mL BMP4 and 10 µM Y-27632 (both from R&D Systems). fl1 medium contains 0.3 ng/mL BMP4 and 10 µM CHIR99021 (R&D Systems). At fl3, a half-medium change was performed with 0.3 ng/mL BMP4, 10 µM CHIR99021, and 10 µM Y-27632. fl5 medium included 0.3 ng/mL BMP4, 10 µM CHIR99021, 10 µM Y-27632, and 2 µM SB431542 (Sigma).

fl7 medium was supplemented with 1 ng/mL BMP4, 2 µM CHIR99021, 10 µM Y-27632, 30 µM SB431542, 0.1 µM retinoic acid (Sigma), and 50 ng/mL human sonic hedgehog (SHH) (R&D Systems). At fl10, the medium contained 3 ng/mL BMP4, 30 µM SB-431542, 50 ng/mL SHH, 10 µM IWR-1 (Sigma), 2 µM SU-5402 (R&D Systems), and 10 µM DAPT (R&D Systems). A half-medium change was performed at fl12 using the same composition as fl10. At fl15, the medium included 2 µM SB-431542, 50 ng/mL SHH, 1 µM IWR-1, 1 µM SU-5402, and 10 µM DAPT.

NT^+^ APLCs were FACS sorted at fl18. A total of 20,000 NT^+^ cells were centrifuged at 210 g for 4 min to promote aggregation in V-bottom 96-well low-cell-binding plates (Greiner Bio-One) using MK10 medium. The plate was incubated at 37 °C under a 5% COL normoxic condition. After 24 hours, the aggregates were transferred to 30 µL of Matrigel on ice, then plated in a pre-warmed 24-well plate (Corning). The plate was inverted and incubated at 37 °C for 10 min to allow Matrigel dome formation before being returned to its original position. A total of 450 µL MK10 medium supplemented with 100 ng/mL RSPO3 and 2 µM SB431542 was added per well, and the medium was changed every three days. NT^+^CD10^+^ DZLCs are FACS sorted between mt14 and mt21.

For non-reporter hiPSC lines, aggregates at fl18 were directly embedded in Matrigel and cultured with MK10 medium containing 100 ng/mL RSPO3 and 2 µM SB431542. As an alternative to Matrigel culture, DZLCs could be induced by culturing 20,000 APLCs in U-bottom 96-well low-cell-binding plates, with a half-medium change performed every three days.

### Induction of TZLCs and FZLCs from DZLCs

NT^+^CD10^+^ DZLCs were FACS-sorted after 14–21 days of culture in MK10 medium containing 100 ng/mL RSPO3 and 2 µM SB431542. A total of 20,000 DZLCs were centrifuged at 160 g for 4 min to promote cell aggregation in V-bottom 96-well low-cell-binding plates in MK10 medium supplemented with 100 ng/mL RSPO3 and 2 µM SB431542. The aggregates were then incubated at 37 °C under 5% COL in normoxic conditions. After 24 hours, the aggregates were plated in a pre-warmed 24-well plate following the same procedure as described above. The aggregates were then cultured in 450 µL of MK10 medium containing 100 ng/mL RSPO3 and 5 nM ACTH for seven days to induce TZLCs. To further induce FZLCs, TZLCs are washed with MK10 medium and then cultured in MK10 medium supplemented with 5 nM ACTH for an additional seven days. In some experiments, TZLCs and FZLCs were induced using a floating culture system in U-bottom 96-well low-cell-binding plates. Half of the medium was replaced every three days.

### Induction of CapLCs

A whole organoid at fl18 was transferred to Matrigel following the same procedure as used for DZLC induction and cultured in MK10 medium without cytokines. The medium was changed every three days. NT^-^RG^+^ CapLCs were FACS-sorted at mt21.

### Transplantation of DZLCs or APLCs encapsulated with CapLCs

Male NOD-*Prkdc^em26Cd52^Il2rg^em26Cd22^*/NjuCrl (NCG) mice, 6–8 weeks (Charles River), were used for transplantation. The mice were housed under a 12:12 hours light-dark cycle at a temperature of 22–24°C and a humidity level of 40–60%. All surgical procedures were performed aseptically in accordance with IACUC guidelines for rodent survival surgery.

Encapsulation was conducted 48 hours after the re-aggregation of FACS-sorted DZLCs and 24 hours after the re-aggregation of FACS-sorted APLCs. First, 10,000 RG^+^ CapLCs were centrifugated in MK10 medium at 210 g for 2 min in V-bottom 96-well low-cell-binding plates. Then, aggregates consisting of 20,000 NT^+^CD10^+^ DZLCs or NT^+^ APLCs were transferred to wells containing the CapLCs. Finally, after additional 10,000 RG^+^ CapLCs were added to wells, the plates were centrifugated at 210 g for 2 min. 48-72 hours after the encapsulation, organoids were transplanted under the kidney capsule of NCG mice that received hemi-adrenalectomy.

### Lentivirus packaging and transduction to APLCs

sgRNAs targeting exon 3 of *CTNNB1* were designed using the Molecular Biology CRISPR design tool (Benchling). The sgRNA sequences were 5’-TGATTTGATGGAGTTGGACA-3’ and 5’-CAACAGTCTTACCTGGACTC-3’, which were cloned into a linearized LentiCRISPRv2GFP vector (a gift from David Feldser, Addgene plasmid #82416; http://n2t.net/addgene:82416; RRID:Addgene_82416).

For lentivirus packaging, the target vector was co-transfected into 293T cells alongside pMD2.G (a gift from Didier Trono, Addgene plasmid #12259; http://n2t.net/addgene:12259; RRID:Addgene_12259) and psPAX2 (a gift from Didier Trono, Addgene plasmid #12260; http://n2t.net/addgene:12260; RRID:Addgene_12260) lentiviral packaging plasmids using PEI MAX (Polysciences, 24765-1). As a control, LentiCRISPRv2GFP without sgRNAs was also packaged.

Lentiviral particles were collected 48 and 72 hours post-transfection, concentrated 300-fold using a Sorvall AH-629 centrifuge rotor (Thermo Scientific, 54284) and Thinwall Ultra Clear Tubes (Beckman Coulter, 344058) at 23,000 rpm for 2 hours. The viral particles were then filtered using Costar Spin-X Centrifuge Tube Filters (Corning, 8160) and added to MK10 medium containing 20,000 FACS-sorted APLCs in V-bottom 96-well low-cell-binding plates. The cells were incubated at 37 °C under 5% COL normoxic conditions for 1 hour, centrifuged at 210 g for 4 min to promote aggregation, and further cultured under the same conditions. After 16 hours, the aggregates were washed with MK10 medium, transferred to Matrigel, and cultured under DZLC induction conditions as described above.

### Fluorescence-activated cell sorting (FACS)

Aggregates were dissociated using 0.1% trypsin/EDTA for 10–15 min at 37 °C, with periodic pipetting to aid dissociation. The reaction was quenched by adding an equal volume of fetal bovine serum (FBS), and the cells were resuspended in FACS buffer (0.1% bovine serum albumin [BSA] in PBS). Before sorting or analysis, cell suspensions were passed through 70-µm nylon cell strainers (Thermo Fisher Scientific) to remove clumps. For cell surface marker analysis and sorting, dissociated cells were stained with BV421-conjugated anti-human CD10 (BioLegend), PE-conjugated anti-human CD10 (BioLegend), FITC-conjugated anti-human CD34 (BioLegend), or BV421-conjugated anti-human CD140a (PDGFRA) (BD Biosciences). Cells stained with surface markers and those expressing NT, RG, or HG were sorted using FACSAria Fusion (BD Biosciences) and collected in 1.5-mL Eppendorf tubes or 5-mL polypropylene round-bottom tubes (Corning). FACS data were acquired using FACSDiva Software v8.0.2 (BD Biosciences) and analyzed with FlowJo software (BD Biosciences). Statistical significance was determined using one-way ANOVA for repeated measures, followed by Dunnett’s multiple comparison test or a two-tailed Welch’s t-test.

### Quantitative RT-PCR

Cells were pelleted by centrifugation at 250 g for 5 min, then lysed for RNA isolation using the RNeasy Micro Kit (Qiagen). For some assays, cDNA synthesis was performed using Superscript III (Invitrogen) with oligo-dT primers and used for qPCR assays without amplification. In other assays, cDNA synthesis and amplification were carried out using 1 ng of purified total RNA, as previously described ^44^. Target cDNAs were quantified using PowerUp SYBR Green Master Mix (Applied Biosystems) with the StepOnePlus system (Thermo Fisher Scientific). LogL gene expression values were calculated using the ΔCt method, normalized to the average Ct values of *PPIA* and *ARBP*. Statistical significance was determined using one-way ANOVA for repeated measures, followed by Dunnett’s multiple comparison test or a two-tailed Welch’s t-test. The significance levels were as follows: *, p < 0.05; **, p < 0.01; ***, p < 0.001.

### Immunofluorescence analyses on frozen sections

The procedure was performed as previously described ^13^. Briefly, aggregates were fixed with 4% paraformaldehyde (Sigma) in PBS for 2 hours at room temperature, then washed three times with PBS containing 0.2% Tween-20 (PBST) for 15 min. The samples were sequentially immersed in 10% and 30% sucrose (Fisher Scientific) in PBS overnight at 4 °C, embedded in OCT (Fisher Scientific), snap-frozen using liquid nitrogen, and sectioned into 10-µm slices using a cryostat (Leica, CM1800). Sections were placed on Superfrost microscope glass slides (Thermo Fisher Scientific) and stored at -80 °C until further analysis.

Before primary antibody incubation, slides were air-dried, washed three times with PBS, and incubated with a blocking solution (5% normal donkey serum in PBS containing 0.2% Tween-20) for 1 hour. For certain antibodies requiring antigen retrieval, slides were post-fixed with 10% buffered formalin (Fisher Healthcare) for 10 min at room temperature, followed by three 10-min washes with PBS. Antigen retrieval was performed using HistoVT One (Nacalai USA) for 20 min at 70 °C, followed by a 10-min PBS wash at room temperature.

Slides were then incubated with primary antibodies in blocking solution for 2 hours at room temperature, followed by six PBS washes (totaling 2 hours). Secondary antibodies and 1 µg/ml DAPI (Thermo Fisher Scientific) in blocking solution were applied for 50 min at room temperature. In some experiments, live tdTomato signals were detected without antibody staining (Fig. 1M, fig. S7D). After six final PBS washes, slides were mounted in Vectashield mounting medium (Vector Laboratories) and analyzed using confocal laser scanning microscopy (FLUOVIEW FV3000, Evident Scientific). Confocal images were processed with Olympus FV-31S-SW (version 2.6.1.243).

### Immunofluorescence analyses on paraffin sections

For immunofluorescence (IF) analyses of fetal adrenal glands and hiPSC-derived organoids, samples were fixed in 10% buffered formalin (Fisher Healthcare) with gentle rocking overnight at room temperature. After dehydration, tissues were embedded in paraffin, sectioned into 4-µm slices using a microtome (Thermo Scientific Microm™ HM325), and placed on Superfrost microscope glass slides.

Paraffin sections were deparaffinized using xylene, and IF staining was performed as previously described ^13^. Briefly, antigen retrieval was conducted by treating the slides with HistoVT One for 35 min at 90 °C, followed by 15 min at room temperature. Slides were then incubated overnight at 4 °C with primary antibodies in blocking buffer. After six PBS washes, secondary antibodies and 1 µg/ml DAPI (Thermo Fisher Scientific) in blocking solution were applied for 50 min at room temperature. Finally, slides were washed six times with PBS before being mounted in Vectashield mounting medium for confocal laser scanning microscopy.

### In-situ hybridization (ISH) on paraffin sections

ISH analysis on formalin-fixed, paraffin-embedded sections was performed as previously described ^13,26^. Briefly, the samples were hybridized using the ViewRNA ISH Tissue Assay Kit (Thermo Fisher Scientific) with gene-specific probe sets for human *RSPO3* (#VA1-13645-VT). Probe sets against Bacillus S. *dapB* (#VF1-11712-VT) served as a negative control.

Deparaffinized sections were treated with a pretreatment solution for 12 min at 90–95 °C, followed by incubation in a protease solution for 6 min and 30 seconds at 40 °C. After fixation in a 10% formaldehyde neutral buffer solution for 5 min, sections were hybridized with the ViewRNA type 1 probe set for 2 hours at 40 °C. This was followed by sequential incubations with the preamplifier probe (25 min at 40 °C), the amplifier probe (15 min at 40 °C), and the Label Probe 1-AP (15 min at 40 °C). Next, sections were treated with an AP enhancer for 5 min at room temperature, followed by signal development using FastRed Substrate Solution for 1 hour at room temperature. Finally, slides were counterstained with DAPI (1 mg/mL) in PBS for 1 hour and mounted in Vectashield mounting medium for confocal laser scanning microscopy.

### Transmission electron microscopy

Tissues for electron microscopy were fixed overnight at 4°C in 2.5% glutaraldehyde and 2.0% paraformaldehyde in 0.1 M sodium cacodylate buffer (pH 7.4). After buffer washes, samples were post-fixed in 2.0% osmium tetroxide for 1 hour at room temperature, then rinsed in dHLO before en bloc staining with 2% uranyl acetate. Tissues were dehydrated through a graded ethanol series, infiltrated, and embedded in EMbed-812 (Electron Microscopy Sciences, Fort Washington, PA). Thin sections were stained with uranyl acetate and lead citrate and examined using a JEOL 1010 electron microscope equipped with a Hamamatsu digital camera and AMT Advantage NanoSprint500 software.

### Enzyme-linked immunosorbent assay (ELISA)

To measure RSPO3 production from whole organoids cultured in MK10 medium under cytokine-free conditions, organoids were derived from 232B6 (*RSPO3* KO) and 2125 (wild-type parental control) hiPSCs. Culture supernatants were collected at mt21, 72 hours after the last medium change. RSPO3 concentration was determined using an ELISA kit (Abnova, 89335201) according to instructions. Data are presented as means ± standard deviation (n = 4). P-values were calculated using a two-tailed Welch’s t-test for statistical comparison. *, p < 0.05; **, p < 0.01; ***, p < 0.001.

To measure steroid hormones in culture supernatants, the culture medium for aggregates was refreshed three days prior to collection for steroid hormone measurement. After 72 hours of incubation, supernatants were collected and stored at –20L°C until analyses. Steroid hormone levels, including DHEA, DHEA-S, and cortisol, were measured using commercially available clinical-grade ELISA kits (DRG International: DHEA [EIA3415], DHEA-S [EIA1562], and Cortisol [EIA-1887R]), following the manufacturer’s instructions. Optical density (OD) at 450 nm (OD450) was determined using a BioTek Epoch microplate reader (Agilent) and analyzed with BioTek Gen6 software (Agilent). Statistical significance was assessed using one-way ANOVA for repeated measures followed by Dunnett’s multiple comparison test or a two-tailed Welch’s t-test.

### Steroid quantification by liquid chromatography-tandem mass spectometry (LC-MS/MS)

Steroid quantitation was performed based on previously described methods with modifications ^45,46^. For Δ4 steroid measurement, a mixture of 0.1 mL culture medium or mouse serum, 0.1 mL deionized water (dHLO), and 0.1 mL internal standards at known concentrations was loaded onto supported liquid extraction (SLE) columns (Isolute; Biotage, Charlotte, NC). After 5 min, the columns were washed twice with 0.9 mL methyl-tert-butyl ether (MTBE), and pressure was applied for 30 sec to complete elution. The eluatesWere dried in a centrifugal vacuum, resuspended in 0.2 mL 40:60 methanol:water, and loaded into the autosampler.

For Δ5 steroid measurement, a mixture of 0.1 mL culture medium or mouse serum and 0.1 mL internal standards in acetonitrile was deproteinized with 0.2 mL methanol and agitated for 10 seconds. After incubation at room temperature for 5 minutes, samples were centrifuged for 5 min, and the supernatant was transferred to an autosampler vial, mixed with 0.4 mL water, and extracted twice with 0.8 mL MTBE. The combined organic extracts (upper layers) were dried in a centrifugal vacuum and derivatized by agitation for 10 sec with 0.1 mL reagent containing 2-picolinic acid, 2-methyl-6-nitrobenzoic anhydride, and 4-(dimethylamino) pyridine (25 mg, 20 mg, and 10 mg per 1 mL of acetonitrile, respectively). The mixture was incubated at room temperature for 90 min in an orbital shaker. The reaction was quenched with 0.4 mL water, and a 0.4 mL aliquot was loaded onto an Isolute SLE column. After equilibration under gentle pressure for 5 min, the column was washed twice with 0.9 mL MTBE, followed by 30 sec of applied pressure to complete elution. The elutes were dried in a centrifugal vacuum, resuspended in 0.4 mL 50:50 methanol:water, transferred to injection vials, and loaded into the autosampler.

For Δ5 steroid sulfate measurement, a 10 µL aliquot of culture medium or mouse serum was diluted with 290 µL water, mixed with 100 µL internal standards, and loaded onto an Isolute SLE column. After equilibration under gentle pressure for 5 minutes, 1.4 mL 1:1 (v/v) chloroform/2-butanol was added and allowed to permeate the column for 10 minutes. After a 5-minute equilibration, gentle pressure was applied to expel the solvent, which was then collected. The solvent was dried for at least 1 hour in a centrifugal vacuum preheated to 55°C. The dried residue was resuspended with agitation in 0.2 mL 40:60 methanol:water, transferred to a taper-bottomed injection vial, and analyzed.

Samples (10 µL) were injected and resolved using a two-dimensional chromatography method with a C4, 10 × 3 mm column on an Agilent 1260 binary pump HPLC system, followed by a Kinetex 150 × 2.1 mm, 2.6 µm particle-size biphenyl column on an Agilent 1290 binary pump UPLC system, using gradient elution with 0.2 mmol/L ammonium fluoride in water and methanol. The column effluent was directed into an Agilent 6490 triple quadrupole tandem mass spectrometer using electrospray ionization in negative ion mode, and data were acquired using MRM mode.

### Bulk RNA-seq library preparation

Fetal adrenal glands, the heart, hiPSCs, and hiPSC-derived adrenal organoids were used for bulk RNA sequencing. Fetal tissues were dissociated using a Multi Tissue Dissociation Kit 1 (Miltenyi Biotec) according to the manufacturer’s instructions. Briefly, tissues were minced into fragments smaller than 1 mm using scissors, collected in 15-mL Falcon tubes, and mixed with a dissociation solution (enzyme mix in serum-free DMEM). The samples were then incubated at 37 °C in a water bath for 30–40 min, with pipetting every 10 min. Following incubation, FBS was added to 15% of the total volume, and cell suspensions were strained through 70-µm cell strainers. Cells were washed once in PBS by centrifugation at 300 g for 5 min and treated with Red Blood Cell Lysis Solution (Miltenyi Biotec) for 4 min at room temperature. After two washes with FACS buffer, cells were centrifuged at 200 g for 5 min and resuspended in FACS buffer before counting via trypan blue exclusion.

For select samples (fetal adrenal glands at 15, 17, and 19 weeks post-fertilization), cells were stained with BV421-anti-PDGFRA, FITC-anti-CD34, PE-anti-CD10 antibodies. CD34-negative CD10-positive cells were sorted as DZ cells, while PDGFRA-, CD34-, and CD10-negative cells were sorted as TZ/FZ cells using a FACSAria Fusion sorter. Cells were pelleted by centrifugation at 250 g for 5 min, snap-frozen in liquid nitrogen, and stored at –80 °C until RNA isolation. Dissociation of hiPSC-derived aggregates was performed as described previously ^26^. RNA was extracted using the RNeasy Micro Kit (Qiagen), and its quality was assessed with the High Sensitivity RNA ScreenTape on an Agilent 4200 TapeStation. cDNA library preparation was performed using the Clontech SMART-Seq mRNA HTLP kit (Takara Bio, 634792) following the manufacturer’s protocol. AMPure XP beads were used for library cleanup. Sequencing was conducted on an Illumina NextSeq 2000 platform, generating 120-base pair reads according to the manufacturer’s protocol.

### Mapping reads and data analysis for bulk RNA-seq

Raw sequencing data were demultiplexed into Fastq files using bcl2fastq2 (v2.20.0.422). Barcode and adaptor sequences were trimmed with Trimmomatic (v0.32). The processed Fastq files were then aligned to the UCSC human reference genome (GRCh38) using STAR (v2.7.1a). A raw gene count table was generated with featureCounts. Differentially expressed genes (DEGs) were analyzed using edgeR (v3.36.0), applying thresholds of log2 fold change > 1, *p*-value < 0.05, and FDR < 0.05. Log2 (RPKM+1) values were used for scatterplot visualization.

### 10x Genomics single-cell RNA-seq library preparation

Samples were dissociated into single cells as previously described, resuspended in 0.1% BSA in PBS, and counted ^26^. Viability exceeded 80% for all samples, as determined by trypan blue staining. Encapsulation was carried out using a 10x Genomics Chromium Controller following the manufacturer’s protocol. Cells were loaded into Chromium microfluidic chips with the Chromium Single Cell 3’ Reagent Kit (v3.1 chemistry) to generate single-cell Gel Bead emulsions (GEMs). Reverse transcription was performed using a C1000 Touch Thermal Cycler with a 96-Deep Well Reaction Module (Bio-Rad). Subsequent cDNA amplification and library construction were conducted according to the manufacturer’s protocol. Libraries were sequenced using a 150-cycle paired-end sequencing protocol on a NextSeq 2000 instrument.

### Read Mapping of scRNA-seq data and downstream analysis

Sequence data was demultiplexed and fastqs produced using Cell Ranger (v.7.1.0). For samples containing only human cells, reads were mapped to the GRCh38 human reference. For human samples transplanted into mice, samples were also independently mapped to the GRCm38 mouse reference. This produced two separate alignments for each cell, one for the human reference and another for the mouse reference. Both sets of UMI counts were read into R (v.4.4.1) and secondary analysis was performed using Seurat (v.5.1.0) ^47^. UMI count tables were first loaded into R by using the Read10x function, and Seurat objects were built from each sample, one per alignment. The sum of UMI counts for each alignment was totaled for each cell for both human and mouse counts. The total UMIs for human alignment were then compared with the mouse alignment. Any cell with greater than twofold assignment of UMIs to human than mouse was designated as a human cell. Similarly, any cell with twofold higher mouse counts than human would be designated mouse. Any cell that failed to meet either threshold was labeled “undetermined” and excluded from the analysis. Human (iPSCs and prenatal 7 week adrenal) samples and mouse (two independent mice using adult tissue) samples were used as controls. No human or mouse control cells were assigned to be the wrong species (fig. S7E). In this fashion, all cells with “mouse” or “undetermined” identities were excluded, and only human cells aligned to the human reference were kept for downstream analysis. For each sample a minimum number of genes = 2,000 and maximum genes = 10,000 were set as well as a maximum mitochondrial content of 25% and cells outside those bounds were excluded. A minimum UMI was set on a per-sample basis to exclude low-quality cells by assigning a minimum cutoff based on an inspection of a ranked plot of UMI per sample. Minimum UMI cutoffs were as follows: DZLC-CapLC_tx_d87, APLC-CapLC_tx_d55 – 10^3.5^; fetal_adrenal_17wpf, adult_adrenal_38yo – 10^3.6^; adult_adrenal_50yo, DZLC_d45 – 10^3.7^; fetal_adrenal_6wpf, fetal_adrenal_8wpf1, fetal_adrenal_8wpf2 – 10^3.9^; TZLC, FZLC, APLC-CapLC_tx_d90 – 10^4^. For all other samples a minimum of 10^3.8^ UMI/cell was set. Cell cycle state was assigned using Seurat’s CellCycleScoring function.

Samples were scaled using SCTransform ^48^, regressing “S.Score” and “G2M.Score”. Layers were integrated using Harmony method within Seurat’s IntegrateLayers function ^48^. The first 30 dimensions were used for FindNeighbors, FindClusters and RunUMAP to generate an integrated object containing all cells. Cortical cells expressing *NR5A1* and capsular cells expressing *RSPO3* were subset from this object (fig. S7G). The process of generating a UMAP and then sub-setting any non-cortical non-capsular cells was repeated to remove any contaminating cells from other cell types. For all figures 15,936 *in vitro*, 11,363 transplanted, 11,753 human prenatal and 24,872 human adult cortical cells were used. Additionally, 1,811 transplanted, 4,006 prenatal and 191 adult capsular cells were captured. Clusters were generated using monocle3 ^49^ using resolution=5.2e-6, num_iter=1, random_seed = 122, k = 12, partition_qval=0.05. Pseudotime was generated using monocle’s learn_graph function with ncenter=50, minimal_branch_len=17. Using previously identified marker genes for capsular cells, DZ, TZ and FZ, clusters were assigned to one of these cell types.

DEGs between groups were determined using Wilcoxon Rank Sum test via Seurat’s FindMarkers function. For pseudobulk analysis, mean normalized counts were used by cell type, distance was calculated using the maximum distance between two components and principle components calculated using prcomp. The first three principle components were graphed. Smoothed heatmaps created using ksmooth of values against pseudotime. Gene ontology enrichment was analyzed via DAVID (v2023q4) ^50^.

### Quantification and statistical analysis

Statistical analysis was conducted using Excel, StataSE (v14.2), and R (v4.3.3). The number of biological replicates and the statistical tests performed are specified in the figure legends or elsewhere in the Methods section. Boxplots display the median (center line), upper and lower quartiles (box limits), and 1.5× interquartile range (whiskers). All fluorescence, histological images, and flow cytometric data represent at least two independent experiments with consistent results, unless otherwise indicated in the figures.

## Acknowledgements

We thank L. B. King for carefully reviewing the manuscript and providing insightful comments. We appreciate W. Yang at iPSC Core facility at the University of Pennsylvania for providing human iPSC lines. We thank The Center for Host-Microbial Interactions and the Women’s Health and Clinical Research Center at the University of Pennsylvania for high-throughput sequencing and human sample collection, respectively. We are grateful for Flow cytometry Core at Children’s Hospital of Philadelphia for their support in FACS sorting and analyses.

This work was supported in part by Japan Society for the Promotion of Science Overseas Research Fellowships to M. M., National Institute of Diabetes and Digestive and Kidney Diseases National Institutes of Health Research Grants (1R01DK134493), the Open Philanthropy funds from the Silicon Valley Community Foundation and the Good Ventures Foundation, Quinlivan Family foundation, and IRM translational Project Award from Marda Foundation to K.S.

## Author contributions

K.S conceived and supervised the project and designed the overall experiments. K.S., E.C.W., M.M. wrote the manuscript. M.M. conducted the overall experiments and analyzed the data. E.W. contributed to the analyses of RNA-seq. M.M., T. S. contributed to the culturing and processing of organoids. A.N.L. contributed to xenograft transplantation. J.F.S. contributed to editing the manuscript. D.G.S., R.J.A contributed to the LC-MS/MS analyses and data interpretation.

## Declaration of interests

A. M. M. and K.S. are inventors on a patent covering adrenocortical organoid technology.

## Legends to Supplemental Figures

**Fig. S1. Canonical WNT signaling provided by CapLCs is essential for derivation of DZLCs in vitro**

**(A)** FACS analysis of 211 iPSC-derived organoids cultured under air-liquid interface (ali) or Matrigel (mt) conditions in FZLC medium (DAPT [5 µM], IWR1 [1 µM], SHH [50 ng/mL], SU5402 [2 µM]). Brightfield (BF) and fluorescence (NT, red) images of the organoids at ali21 or mt21 were captured, and the number of NT^+^ cells was quantified by FACS. Data are presented as mean + standard deviation (n = 4). *p < 0.05, **p < 0.01, ***p < 0.001. Scale bar, 250 µm.

**(B)** DHEA and DHEA-S production in the culture supernatant from (A). Culture medium was collected at 72 hours. Data are presented as mean + standard deviation (n = 4). *p < 0.05, **p < 0.01, ***p < 0.001.

**(C)** Schematic depicting CRISPR/Cas9 gene editing of the *RSPO3* locus to establish *RSPO3* knockout (KO) iPSCs (232B6). Black boxes represent exons, and arrows indicate the gRNA targeting regions. Genotyping PCR was performed to confirm the indel (top right). Frameshift mutation confirmation by Sanger sequencing is shown (bottom). WT, wild-type parental 2125 iPSCs.

**(D)** Karyotype (top left) and phase-contrast image (top right) of 232B6 (*RSPO3* KO). A normal karyotype is confirmed. Scale bar, 100 µm. RSPO3 levels in the culture supernatant of whole organoids (bottom right). Culture medium was collected at 72 hours, and RSPO3 concentration was determined by ELISA. Data are presented as mean + standard deviation (n = 4). *p < 0.05, **p < 0.01, ***p < 0.001.

**(E)** IF images of fetal adrenal glands at 5 and 6 weeks of post-fertilization (wpf), stained for CD10 (green), NR5A1 (red), and their merged images with DAPI (gray). Scale bar, 30 µm. AP, adrenal primordium; Ao, aorta; Co, coelom.; D, dorsal; V, ventral; M, medial; L, lateral.

**(F)** IF of the coelomic angles at 6 wpf, stained for CD10 (green), NR5A1 (red), and their merged images with DAPI (gray) (left). In situ hybridization (ISH) on the neighboring section, stained for RSPO3 (red) merged with DAPI (gray). Scale bar, 30 µm. Mes, mesonephros.

**(G)** Schematic illustration of the human *NR5A1* locus and the targeting construct used to generate *NR5A1-3×FLAG-P2A-tdTomato* (*NT*) alleles. Black boxes represent exons. Genotyping PCR was performed to screen for clones bearing *NT* (middle). Clone C12 (designated as SV20-8C12) was identified as monoallelic for *NT*, while the other allele was confirmed to be free of indels by Sanger sequencing. No random integration of the targeting vector (vector) was detected. Phase-contrast image (top right) and karyotype analysis (bottom right) of SV20-8C12 confirming a normal karyotype. Scale bar, 100 µm. Rec., Cre-recombined; Non., non-targeted; WT, wild type; PC, positive control.

**(H)** Schematic illustration of the human *RSPO3* locus and the targeting construct used to generate *RSPO3-P2A-EGFP* alleles (*RG*). SV20-8C12 as described in (G) was used as the parental iPSC line. Black boxes represent exons. Genotyping PCR was performed to screen for clones bearing *RG*. Clone 4 (designated as SV20R2C4) was identified as having biallelic *RG* alleles and showed no random integration of the targeting vector. Phase-contrast image (bottom left) and karyotype analysis (bottom right) of SV20R2C4 confirming a normal karyotype. Floxed, flanked by loxP sites. Scale bar, 100 µm.

**(I)** FACS analysis at the indicated stages of organoids derived from SV20R2C4 hiPSCs cultured under cytokine-free conditions. The number of RGL cells per organoid was quantified by FACS. Data are presented as mean + standard deviation (n = 3). *p < 0.05, **p < 0.01, ***p < 0.001 versus mt5.

**(J)** Heatmap showing the expression of R-spondins across three groups (NT, CapLCs, DN) based on transcriptomic analysis.

**Fig. S2. Reconstitution of the niche allows robust induction of DZLCs in vitro**

**(A)** FACS analysis of organoids cultured with Matrigel (mt14) or floating culture (re-fl14) during RSPO3-mediated DZLC induction from NT^+^ APLCs. The number of CD10^+^ DZLCs was quantified and presented as mean + standard deviation (n = 3).

**(B)** FACS analysis and BF and fluorescence images (NT, red) of 2125-derived organoids at re-fl15, cultured with various R-Spondins (100 ng/mL) for DZLC induction from NTL APLCs. The number of CD10L DZLCs was quantified by FACS. Data are presented as mean + standard deviation (n = 4). *p < 0.05, **p < 0.01, ***p < 0.001 versus RSPO1. Scale bar, 200 µm.

**(C)** FACS analysis of 2125-derived organoids cultured with or without SB431542 (2µM) during DZLC induction by RSPO3 (100 ng/ml). The number of CD10^+^ DZLCs was quantified by FACS. Data are presented as mean + standard deviation (n = 3). *p<0.05, **p<0.01, ***p<0.001.

**(D)** FACS analysis and BF and fluorescence images (NT, red) of 2125-derived organoids at mt21, cultured with or without DAPT (10uM) during DZLC induction by RSPO3 and SB431542. The number of CD10L DZLCs was quantified by FACS. Data are presented as mean + standard deviation (n = 3). *p < 0.05, **p < 0.01, ***p < 0.001. Scale bar, 200 µm.

**(E)** FACS analysis of organoids derived from 2125 iPSCs at the indicated stages during DZLC induction. The number of CD10L DZLCs was quantified. Data are presented as mean + standard deviation (n = 3). *p < 0.05, **p < 0.01, ***p < 0.001 versus fl18.

**(F)** qPCR analysis of key DZ markers at indicated stages. Cells were derived from 2125 iPSCs. *p<0.05, **p<0.01, ***p<0.001 versus fl18.

**(G)** FACS analysis of organoids derived from 2125 hiPSCs, treated with RSPO3 (100 ng/ml) and SB431542 (2 µM) for 21 days, following FACS sorting of NTL APLCs at fl15, fl18, and fl21. The proportion of CD10L cells was quantified. Data are presented as mean + standard deviation (n = 3). *p < 0.05, **p < 0.01, ***p < 0.001 versus fl15.

**(H)** IF images of DZLCs at mt21 and DZ and FZ at 17 wpf, stained for LEF1, KI67, STAR, CYP11A1, CYP17A1, SULT2A1, and POR (green), along with NR5A1 (red), with merged images shown. Scale bar, 30 µm.

**(I)** Schematic illustration of the human *NR5A1* locus and targeting construct to generate *NR5A1-p2A-tdTomato* alleles (*NT*) (top). Black boxes indicate exons. Genotyping PCR was performed to screen for clones bearing *NT* (second panel). Clone 3 (designated as 2123) was identified as monoallelic for *NT*, while the other allele remained free of indels, as confirmed by Sanger sequencing (third panel). No random integration of the targeting vector was observed. Phase-contrast image and karyotype analysis of 2123 (bottom) confirm a normal karyotype. Scale bar, 100 µm.

**(J)** FACS histograms of DZLCs at mt21 derived from 2123 (NT), 211 (WT1-2A-EGFP [WG]; NT), SV20 (non-reporter) hiPSC lines.

**(K)** IF images of DZLCs at mt21 derived from indicated iPSC lines, stained for CD10, NOV, LEF1 (green), NR5A1(red), and their merged images with DAPI (gray). Scale bar, 10 µm.

**Fig. S3. Role of WNT signaling in DZLC induction**

**(A)** Lentiviral transduction of 2125-derived APLCs at fl18–19 with either a *CTNNB1*-gRNA vector or a backbone vector lacking gRNA, followed by DZLC induction (top left). FACS analysis of organoids at mt14 for NT and EGFP expression (top right). CD10 expression in the indicated cell fractions is shown in histograms (bottom left). The percentage and number of CD10L cells were quantified. Data are presented as mean + standard deviation (n = 3). *p < 0.05, **p < 0.01, ***p < 0.001 versus EGFPL cells in the backbone-only group.

**(B)** Effect of the porcupine inhibitor LGK974 (1 µM) on DZLC induction. DMSO was used as the vehicle control. Cells were derived from 2125 iPSCs. The percentage of CD10L cells (top) and absolute cell numbers per organoid (bottom right) were quantified by FACS. Data represent mean + standard deviation (n = 3). *p < 0.05, **p < 0.01, ***p < 0.001. Bright-field (BF) and fluorescence images (NT, red) of organoids at mt21 (bottom left). Scale bar, 200 µm.

**(C)** Schematic illustration of the human *WNT4* locus and targeting region to generate *WNT4* KO iPSCs (top left). Black boxes represent exons, and black arrows indicate the gRNA targeting region. Genotyping PCR confirmed biallelic indels in clone 11 (designated as 237B11) (middle), which were further validated by Sanger sequencing (bottom). Phase-contrast image of 237B11 iPSCs (top right). Scale bar, 100 µm.

**(D)** Generation of *WNT2B* KO iPSCs. Genotyping PCR confirmed biallelic indels in clone 9 (designated as 238A9), which was further validated by Sanger sequencing. Phase-contrast image of 238A9 (top right). Scale bar, 100 µm.

**(E)** Karyotype analysis of 237B11 and 238A9. A normal karyotype was confirmed for both clones.

**(F)** Generation of *ZNRF3* KO iPSCs. Genotyping PCR confirmed biallelic indels in clone 3 (designated as ZT1C3), which was further validated by Sanger sequence. Phase-contrast image of ZT1C3 (top right). Scale bar, 100 µm.

**(G)** Karyotype analysis of ZT1C3. A normal karyotype was confirmed.

**(H)** FACS analysis of *ZNRF3* KO organoids. NT^+^ APLCs were cultured under no cytokine conditions ± LGK974 (1 µM). DMSO was used as vehicle control. The number of CD10L DZLCs was quantified. Data represent mean + standard deviation (n = 3). *p < 0.05, **p < 0.01, ***p < 0.001.

**(I)** Previously identified DEGs of human DZ cells (Table S2) are highlighted in red in the scatter plot in Fig. 2J.

**(J)** Scatter plot comparing average gene expression between DZLCs and FZLCs. DEGs (log2-fold change > 1, FDR < 0.05, log2[RPKM + 1] > 1) are highlighted in color. Representative DEGs and their GO enrichments are shown on the right.

**Fig. S4. Coordinated action of canonical WNT signaling and ACTH mediates transdifferentiation of human prenatal adrenal cortex**

**(A)** IF images of fetal adrenal glands at 18 weeks post-fertilization (wpf) for CD10, SULT2A1 (green), and HSD3B2 (red), with merged images including DAPI (gray). Dotted lines denote DZ/TZ and TZ/FZ boundaries. Scale bar, 30 µm.

**(B)** High magnification of IF images of TZ cells at 18 wpf. Double positive cells are indicated by arrows. Scale bar, 10 µm.

**(C)** IF images of fetal adrenal glands at 18 wpf for LEF1 (green) and HSD3B2 (red), with merged images including DAPI (gray). A high magnification merged image of the indicated region is shown in the inset. Scale bar, 50 µm.

**(D)** IF images of DZLCs and TZLCs treated with RSPO3 (100 ng/ml) and the indicated concentrations of ACTH (0.2, 1, 5 nM) (left). Scale bar, 30 µm. Quantification of CYP17A1, HSD3B2, and double-positive (DP) cells (right). Data are presented as mean + standard deviation (n = 3). *p < 0.05, **p < 0.01, ***p < 0.001 versus DZLCs.

**(E)** IF images of organoids induced into TZLCs in the presence of PKA inhibitors, PKI(5-24) (100 nM) or KT5720 (1 uM). TZLCs were induced by RSPO3 and ACTH for 7 days following FACS-sorting of DZLCs at mt15. Scale bar, 30 µm. Quantification of CYP17A1, HSD3B2, and double-positive (DP) cells (right) by IF. Data are presented as mean + standard deviation (n = 3). *p < 0.05, **p < 0.01, ***p < 0.001 versus Vehicle.

**(F)** Cortisol production from organoids as described in (D) and (E). Culture supernatants were collected at 72 hours, and cortisol concentration was determined by LC-MS. Data are presented as mean + standard deviation (n = 3, except ACTH 5 µM group and Vehicle control in PKA inhibitor experiments, n = 2). *p < 0.05, **p < 0.01, ***p < 0.001 versus DZLCs or Vehicle.

**(G)** IF images of DZLCs, TZLCs, and FZLCs (all derived from 2125 iPSCs), stained for the indicated markers. Scale bar, 30 µm.

**(H)** IF quantification of the data described in (G). Data are presented as mean + standard deviation (n = 3). *p < 0.05, **p < 0.01, ***p < 0.001 versus DZLCs.

**(I)** IF images of indicated organoids derived from SV20 non-reporter iPSCs. Scale bar, 30 µm.

**(J)** Schematic illustration of the human *HSD3B2* locus and targeting construct for generating *HSD3B2-p2A-EGFP* alleles (*HG*). Genotyping PCR to screen for clones bearing *HG*. Note that clone 29 (designated as MH2C29) bears monoallelic *HG*, whereas the other allele is free of indels, which was further confirmed by Sanger sequencing. Random integration of the targeting vector is not observed. Phase-contrast image and karyotype analysis of MH2C29. A normal karyotype is confirmed. Scale bar, 100 µm.

**Fig. S5. RSPO3 and ACTH mediate transdifferentiation of long-term cultured DZLCs in vitro and DZ cells ex vivo**

**(A)** FACS plot of DZLCs (derived from 2125 iPSCs) at mt50 cultured with DZ medium.

**(B)** BF and fluorescence (NT, red) images of DZLCs at indicated stages. Scale bar, 200 µm.

**(C)** IF images of DZLCs at mt50. Scale bar, 30 µm.

(**D, E**) IF images of DZLCs at mt50, TZLCs, and FZLCs induced from DZLCs at mt50 (left), and their quantification. Data are presented as mean + standard deviation (n = 4). *p < 0.05, **p < 0.01, ***p < 0.001 versus DZLCs. Bar, 30 µm.

**(F)** Cortisol, DHEA, and DHEA-S production as in (D, E). Culture medium was collected at 72 hours, and concentrations were measured by LC-MS. Data are presented as mean + standard deviation (n = 4). *p < 0.05, **p < 0.01, ***p < 0.001 versus DZLCs.

**(G)** Gating strategy for FACS-sorting of DZ and TZ/FZ cells from fetal adrenal glands. PDGFRA^−^CD34^−^CD10^+^ cells were collected as DZ cells, while PDGFRA^−^CD34^−^CD10^−^ cells were collected as TZ/FZ cells.

**(H)** Schematic showing FACS sorting of fetal adrenal cells followed by organoid culture in DZ medium.

**(I)** qPCR analysis of freshly isolated DZ and TZ/FZ cells from fetal adrenal glands at 15–17 wpf, as described in (H). DZ markers (first and second panels) and TZ/FZ markers (third panel) were assessed. Data are presented as mean ± standard deviation (n = 2). *p < 0.05, **p < 0.01, ***p < 0.001.

**(J)** IF images of indicated organoids derived from fetal adrenal glands at 17 wpf. Reaggregated organoids were cultured in DZ medium until mt14 following FACS-sorting as in (H). Scale bar, 30 µm.

**(K)** Schematic illustrating the transdifferentiation process of DZ cells ex vivo. FACS-sorted CD10^+^ DZ cells from fetal adrenal glands at 17 wpf were embedded in Matrigel and cultured with RSPO3 and SB431542 for 3 days (exDZ), followed by 7 days in TZ medium (RSPO3 and ACTH, exTZ), and an additional 7days in FZ medium (ACTH only, exFZ).

**(L)** IF images of exDZ, exTZ, and exFZ organoids, stained for indicated markers. Scale bar, 30 µm.

**(M)** IF quantification of samples described in (L). Mean + standard deviation (n = 4). *p<0.05, **p<0.01, ***p<0.001 versus exDZ.

**(N)** Cortisol and DHEA-S production from exDZ, exTZ, and exFZ organoids. Culture supernatants were collected at 72 h, and concentrations were determined by LC-MS. Data are presented as mean + standard deviation (n = 3). *p < 0.05, **p < 0.01, ***p < 0.001 versus exDZ.

**(O)** Boxplots of nucleus and cell size of exDZ, exTZ, and exFZ quantified by IF (n = 100). *p<0.05, **p<0.01,***p<0.001 versus exDZ.

**Fig. S6. Critical role of *NR0B1* and *GATA6* in DZLC differentiation**

**(A)** *NR0B1* in the indicated samples. iPSCs, fl9/10, fl14/15, APLCs, DZLCs, and FZLCs were derived from 211 and 2125 hiPSCs (n = 4). DZ and FZ cells were isolated from fetal adrenal glands at 15, 17, and 19 wpf (n = 3).

**(B)** Schematic illustration of human *NR0B1* locus and targeting region to generate *NR0B1* KO iPSCs. Black arrowheads indicate CRISPR gRNA targeting regions.

**(C)** Genotyping PCR showing indels in 231C3 iPSCs (female, top left), further confirmed by Sanger sequence (bottom). A phase-contrast image of 231C3 (top right). Scale bar, 200 µm.

**(D)** Genotyping PCR showing indels in 242-12 iPSCs (male, top left), further confirmed by Sanger sequence. A phase-contrast image of 242-12 (bottom left). Scale bar, 200 µm.

**(E)** Karyotype analysis of 231C3 and 242-12, both showing a normal karyotype.

**(F)** FACS analysis of organoids at fl18 and mt18. Organoids were cultured in DZ medium until mt18 following FACS-sorting of APLCs at fl18. Organoids were derived from SV20-8C12 (WT) and 242-12 (*NR0B1* KO#2). The number of CD10^+^ cells was quantified at mt18. Mean + standard deviation (n = 4). *p<0.05, **p<0.01,***p<0.001. BF and fluorescence (NT, red) images of the organoids at mt18 are shown at the bottom left. Scale bar, 200 µm.

**(G)** Scatter plot comparing the average gene expression of NT^+^ APLCs at fl18 between 2125 (WT) and 231C3 (*NR0B1* KO) lines. DEGs (log-fold change > 1, FDR < 0.05, log2[RPKM+1] > 1) are highlighted in colors. Gray dots represent non-significant genes. Representative DEGs and their associated GO enrichments are shown at the bottom.

**(H)** Venn diagram showing down-regulated DEGs in *NR5A1* KO organoids at fl18 and up-regulated DEGs in *NR0B1* KO organoids at fl18, both compared with wild-type organoids at fl18. Up- or down-regulated DEGs were identified by bulk transcriptome analysis of FACS-sorted NT^+^ cells (log2-fold change > 1 and FDR < 0.05). Representative overlapping genes are shown at the bottom.

**(I)** IF images of organoids derived from SV20-8C (WT) or 242-12 (NR0B1 KO#2) cultured in DZ medium (RSPO3 and SB431542) until mt18, stained for indicated markers. Scale bar, 20 µm.

**(J)** DHEA and DHEA-S production from organoids in (I). Culture supernatants were collected at 72 hours. Data are presented as mean + standard deviation (n = 4). *p < 0.05, **p < 0.01, ***p < 0.001.

**(K)** IF images of organoids stained for CYP17A1 (green), HSD3B2 (red), and their merged images with DAPI (gray). Organoids were cultured in TZ medium (RSPO3 and ACTH) for 7 days following FACS-sorting and reaggregation of NT+CD10+ cells at mt21. Cells were derived from 2125 (WT) and 231C3 (NR0B1 KO). Scale bar, 30 µm.

**(L)** FACS analysis of 2125-derived FZLC organoids at the indicated stages, directly induced from APLCs by culturing in the presence of DAPT (5 µM), IWR1 (1 µM), SHH (50 ng/ml), and SU5402 (2 µM), following FACS sorting and reaggregation of NT+ APLCs at fl18. The number of CD10+ cells was quantified.

**(M)** qPCR analysis of indicated FZ markers in APLCs (fl18) and FZLCs (mt21) derived from 2125 iPSCs via direct pathway. *p<0.05, **p<0.01, ***p<0.001.

**(N)** IF images of organoids cultured without cytokines following FACS-sorting of NT+ APLCs. Merged DAPI images (gray) are shown. Scale bar, 20 µm.

**(O)** DHEA and DHEA-S production from organoids in (N). Culture supernatants were collected at 72 hours. Mean + standard deviation (n = 4) *p<0.05, **p<0.01,***p<0.001.

**(P)** FACS analysis of organoids from (N) (left). BF and fluorescence (NT, red) images (right). Scale bar, 200 µm.

**(Q)** Generation of GATA6 KO iPSCs. Sanger sequence confirmed indels (left). Phase-contrast images of GATA6 KO iPSCs, 211e3c7 (right). Bar: 200 µm (right).

**(R)** Karyotype analysis of 211e3c7 showing a normal male karyotype.

**Fig. S7. Reconstruction of functional zonation following transplantation of encapsulated APLCs and DZLCs into NCG mice.**

**(A, B)** Unencapsulated DZLC (A) or APLC (B) organoids derived from 2125 hiPSCs were transplanted under the kidney capsule of NCG mice that received hemi-adrenalectomy. BF images of explants at indicated days following transplantation with insets showing high magnification images of organoids (left). Scale bar, 2 mm. IF images of the organoids merged with DAPI (gray) (right). Scale bar, 50 µm.

**(C)** Encapsulated APLCs transplanted as in (a, b). APLCs were derived from 2125 hiPSCs while CapLCs were derived from SV20R2C4 hiPSCs. BF image of the explant (bottom left) and magnified BF and fluorescence (NT, red) images of organoids at d63 post-transplantation are shown on the bottom right. Scale bar, 2 mm.

**(D)** IF images of encapsulated APLCs at d63 post-transplantation, stained for indicated markers. Merged images with DAPI (gray) are shown on the right. Scale bar, 50 µm.

**(E)** Proportion of cells identified as mouse-derived, human-derived, or of undetermined origin in scRNA-seq data. Control samples include mouse (1,2), and human (adrenal, iPSCs) with known origins.

**(F, G)** Unbiased clustering of all adrenal cells projected onto UMAP plot (left) and annotation of cortical (blue) and capsule (brown) cells (right) according to marker expression (G).

**(H)** Dot plots of key DEG expression by cell type.

**(I)** Heatmap showing dynamic gene expression along pseudotime trajectory. Top 300 genes by percentage difference score > 0.03, log2-fold change > 0.35 and q-value < 0.05 were used. The color indicates the Z-score of log-scaled expression.

**(J)** IF images of encapsulated APLCs at d63 post-transplantation, stained for the indicated markers. Insets show high magnification of the area outlined by white dotted lines. Scale bar, 50 µm. Arrows indicate CYP17A1 and HSD3B2 double-positive cells. Yellow dotted lines denote the border between the capsule and cortex.

**(K)** Serum cortisol levels in mice that received encapsulated APLCs (days 63-90 post-transplantation) and non-transplanted control mice. Cortisol levels were determined by LC-MS. Data are presented as mean + standard deviation (n = 3 for encapsulated APLCs, n = 2 for control) *p<0.05, **p<0.01, ***p<0.001. nd, not detected.

## Supplementary Table

**Table S1**

Differentially expressed genes (DEGs) identified from a pairwise comparison between RG^+^ CapLCs and NT^+^ cortical cells at mt21, related to Fig. 1H.

**Table S2**

Previously identified DEGs for Cap and DZ cells compared with other fetal adrenal cell types, related to Fig. 1I and fig. S3I.

**Table S3**

DEGs identified from a pairwise comparison of DZLCs vs. APLCs, or DZLCs vs. FZLCs, related to Fig. 2J and fig. S3J.

**Table S4**

DEGs identified from a multi-group comparison of iPSCs, PIM (fl10), APLCs (fl15/18), and DZLCs (mt21), related to Fig. 2K.

**Table S5**

DEGs identified from a pairwise comparison of DZLCs vs. TZLCs, or TZLCs vs. FZLCs, related to Fig. 4B, C.

**Table S6**

Transcription factors expressed in DZLCs or DZ cells compared with hiPSCs.

**Table S7**

DEGs identified from a pairwise comparison of *NR0B1* KO (231C3) vs. WT (2125) in the whole NT^+^ fraction at mt21 or NT^+^CD10^+^ fraction at mt21, related to Fig. 5G, H.

**Table S8**

Gene lists upregulated in *NR0B1* KO and down regulated in *NR5A1* KO, related to fig. S6H.

**Table S9**

Primers, antibody, and reagents used in this study

