## Supplementary figures and images for "Reconstitution of adrenocortical functional zonation from human pluripotent stem cells"

### Fig. S1

#### Supplementary Figure 1

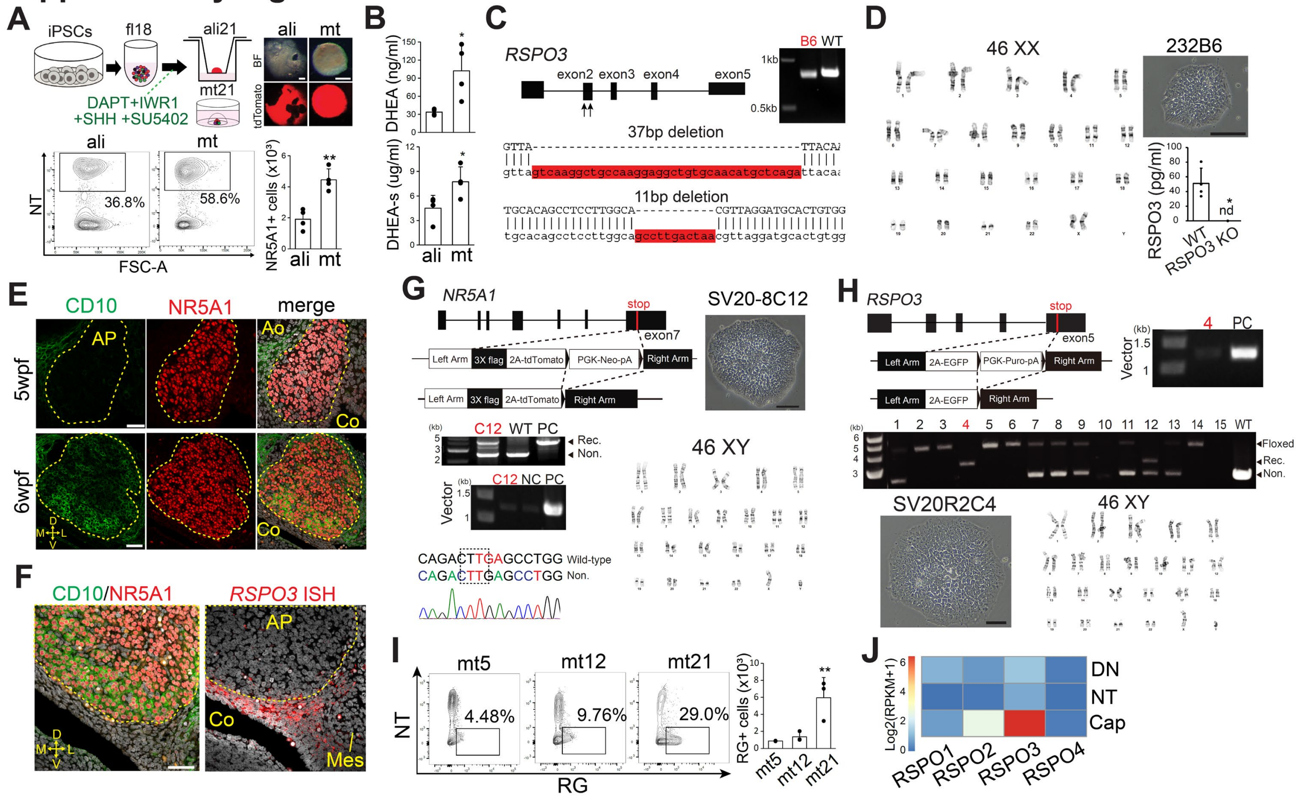

### Fig. S2

#### Supplementary Figure 2

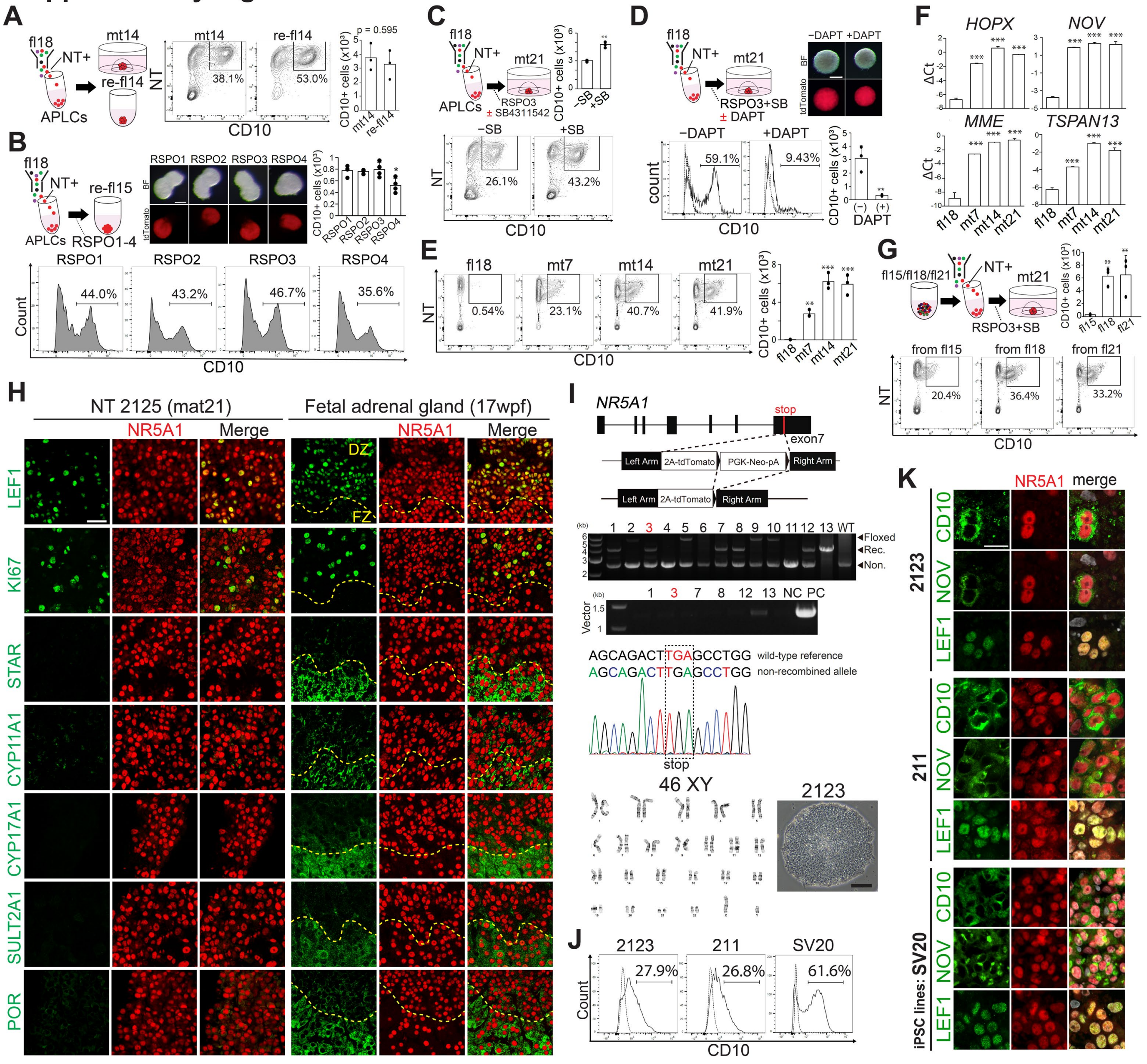

### Fig. S3

#### Supplementary Figure 3

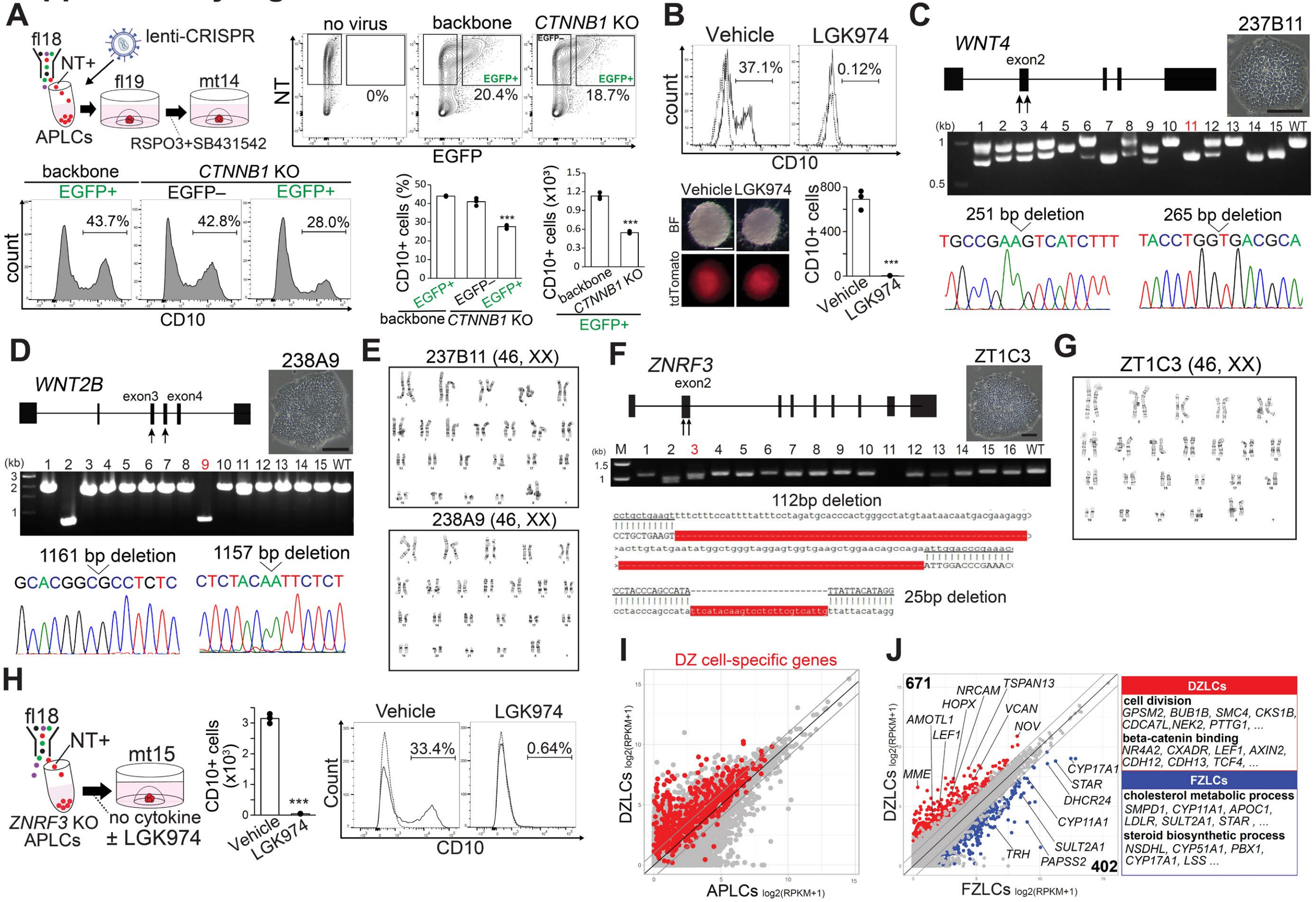

### Fig. S4

#### Supplementary Figure 4

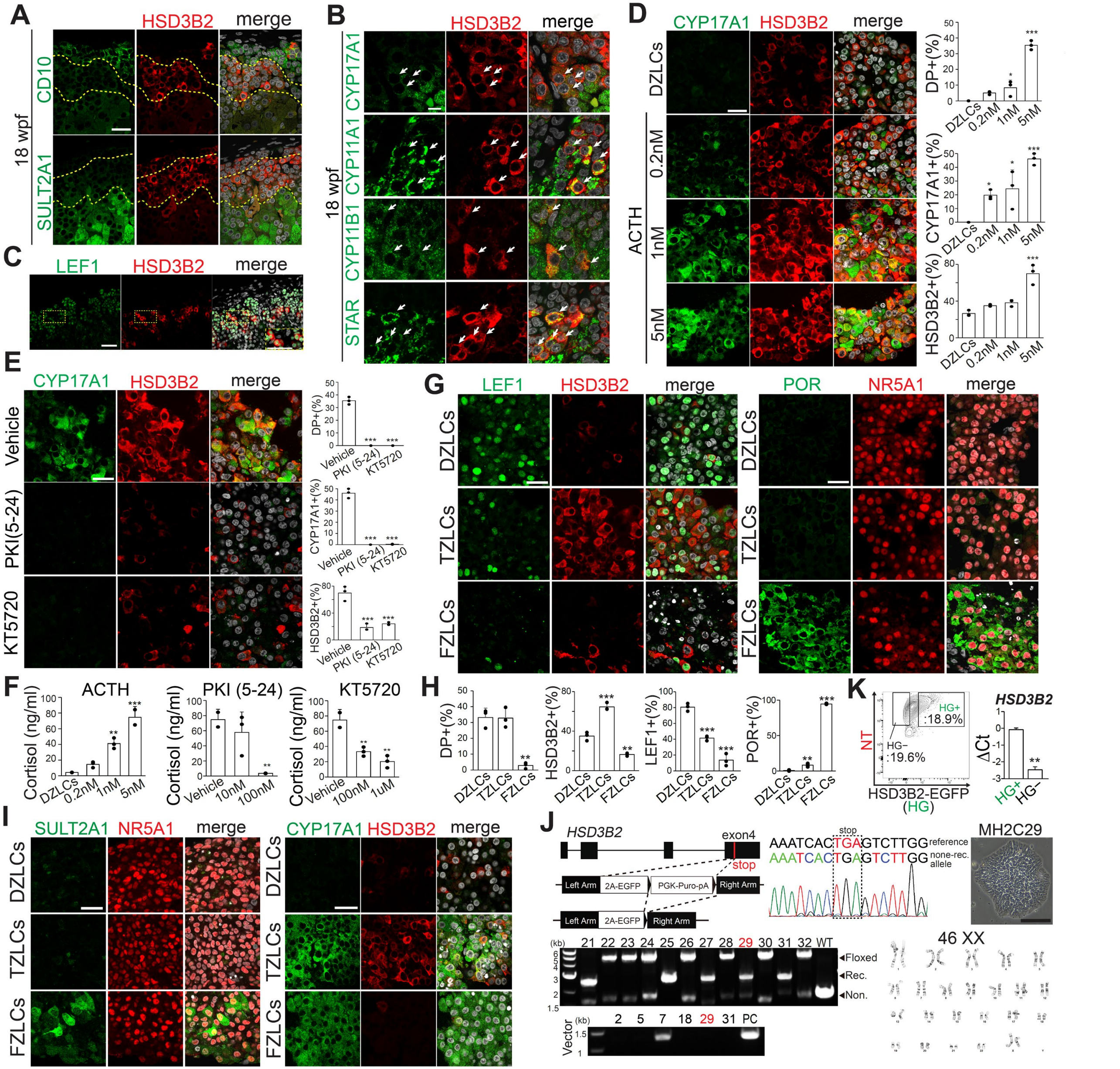

### Fig. S5

#### Supplementary Figure 5

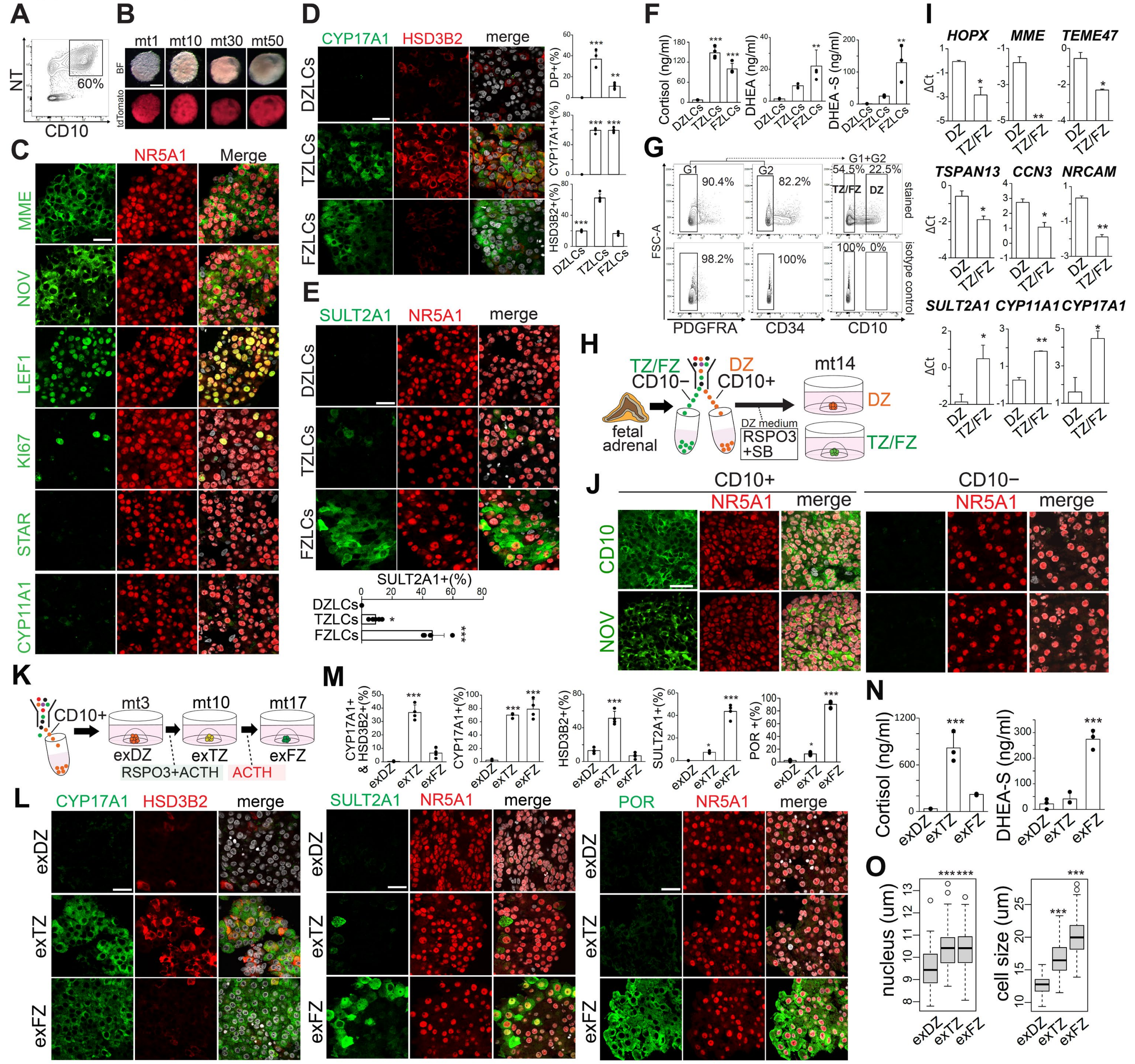

### Fig. S6

#### Supplementary Figure 6

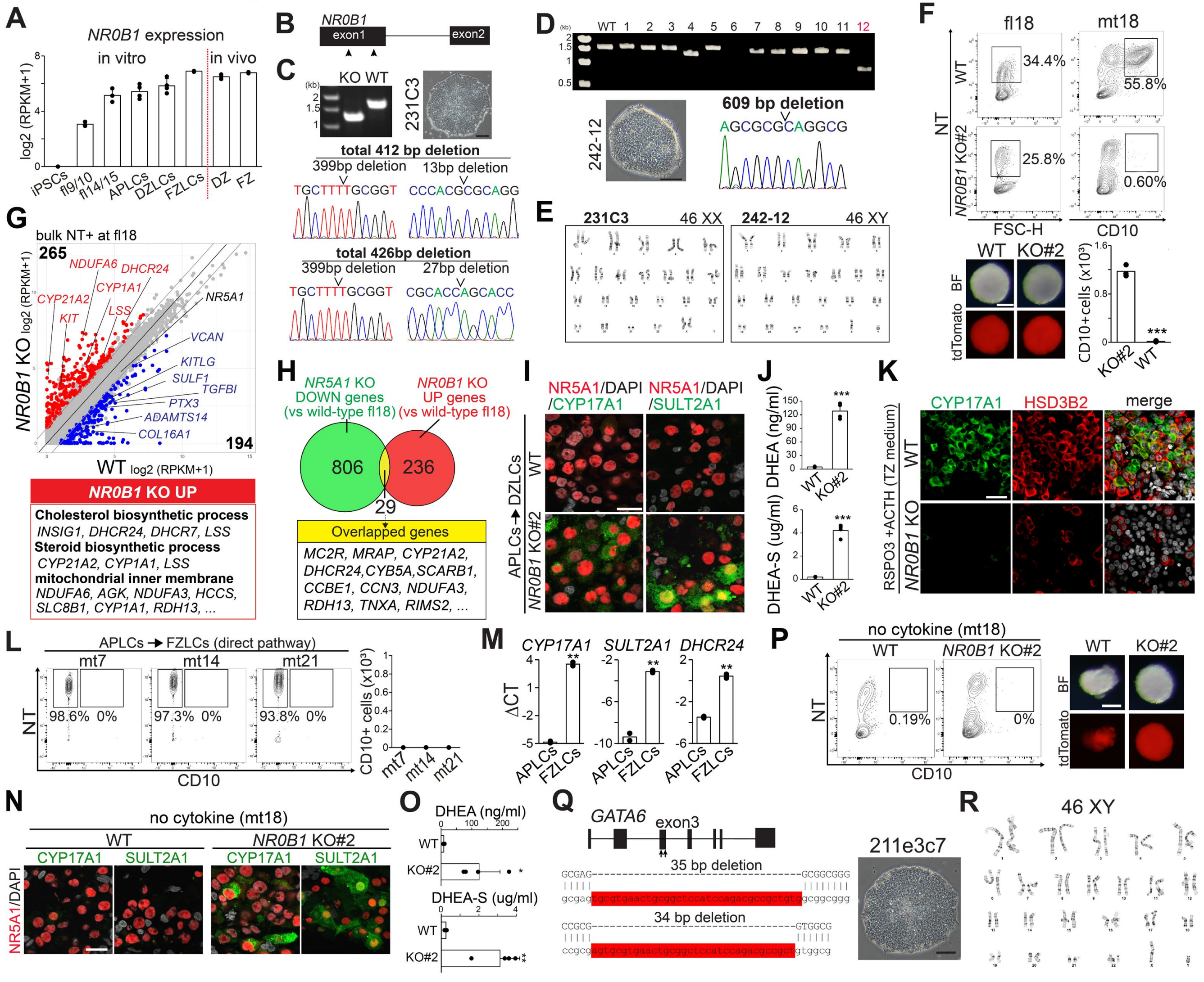

### Fig. S7

##### Supplementary Figure 7

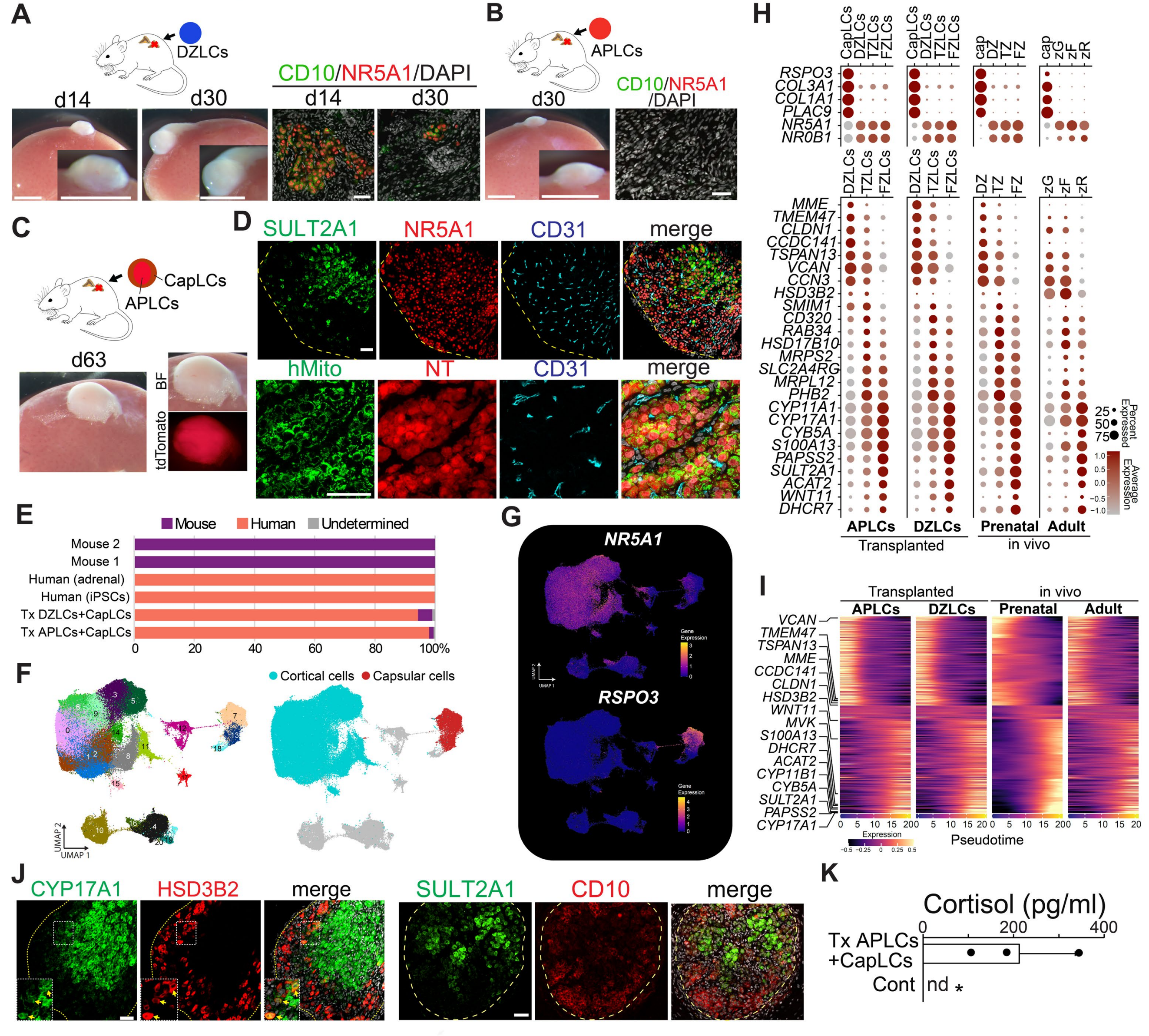
